# Microbial metabolism of methotrexate produces a STAT3 signaling molecule that alleviates gut inflammation

**DOI:** 10.64898/2026.03.11.711168

**Authors:** Juyeong Cho, Rebecca Danner, Woo Sung Kim, Jonathan Williams, Anshit Singh, Jiyoung A. Han, Jonathan G. Van Vranken, Allison S. Walker, Timothy Rhoads, Snehal N. Chaudhari

**Author notes:** Corresponding author: Snehal N. Chaudhari PhD University of Wisconsin-Madison, 433 Babcock Drive, Madison, WI 53706.

## Abstract

Methotrexate (MTX) therapy in inflammatory bowel disease (IBD) is often limited by inter-individual variability in clinical response and adverse effects. Gut microbiota contribute to MTX therapeutic response and toxicity by metabolizing MTX and altering its bioavailability. However, how inflammation alters microbial MTX metabolism and how its metabolites influence the host remain poorly understood. Here, we identify *Clostridium asparagiforme* as a potent and efficient metabolizer of methotrexate, producing deoxyaminopteroic acid (DAMPA) in the distal gastrointestinal tract. We demonstrate that DAMPA preserves mitochondrial integrity by promoting mitophagy in intestinal epithelial cells through mitochondrial STAT3 signaling. DAMPA administration attenuates intestinal inflammation *in vivo*, and improves metabolic dysfunction associated with IBD. Together, these findings reveal an unappreciated role for a gut microbial MTX metabolite in mediating epithelial homeostasis during intestinal inflammation, thus reframing microbial MTX metabolism from passive drug detoxification to active regulation of host mitochondrial and inflammatory homeostasis.

## INTRODUCTION

Methotrexate (MTX), a folate antagonist, is the gold standard therapy for various cancers and inflammatory diseases.^1,2^ MTX suppresses the proliferation of cancer and inflammatory cells by inhibiting Dihydrofolate Reductase (DHFR), a crucial enzyme required for DNA synthesis.^3,4^ Despite its widespread use, MTX therapy is limited by highly variable bioavailability and severe toxicity.^5^ Therapeutic response to MTX is particularly poor in inflammatory bowel disease (IBD) patients, with nearly 75% of individuals failing to achieve clinical efficacy.^6–11^ The most common adverse effects of MTX, including vomiting, diarrhea, and abdominal discomfort, occur in the gastrointestinal tract and can progress to severe complications such as mucosal necrosis, ultimately limiting its clinical use.^12,13^ Moreover, the non-selective cytotoxicity of MTX on rapidly proliferating intestinal epithelial cells can exacerbate gut inflammation.^14^

Inflammation profoundly reshapes gut microbiota composition,^15^ thereby altering bacterial drug metabolism and host responses. In IBD, inflammation-associated microbiota alterations may increase susceptibility to MTX-induced toxicity.^16^ In parallel, MTX can disrupt intestinal barrier integrity and promote bacterial translocation, further amplifying inflammatory responses.^17^ Further, MTX itself can perturb gut microbiota, potentially influencing therapeutic response and immune activation.^18^ However, it remains unclear how inflammation-induced shifts in the gut microbiota modulate MTX metabolism, and subsequent host epithelial and immune responses.

Gut microbiota plays a critical role in shaping MTX efficacy, toxicity, and host immune responses, as differences in gut microbial composition are associated with variability in clinical response to MTX.^19,20^ One mechanism underlying this association is the ability of gut microbiota to enzymatically metabolize MTX, generating the metabolite deoxyaminopteroic acid (DAMPA) through the Carboxypeptidase G2 (CPDG2) enzyme.^21–23^ Previous studies have shown that depletion of gut microbiota exacerbates MTX-induced gastrointestinal damage and promotes pro-inflammatory macrophage accumulation, whereas oral gavage of *Bacteroides fragilis* mitigates these effects through production of immunomodulatory metabolites such as butyrate.^12,24^ These findings establish gut bacteria as both metabolic modifiers of MTX and active regulators of host inflammatory responses.

Despite the growing recognition of microbial MTX metabolism, efforts to identify the specific gut bacterial species responsible for MTX detoxification have been hindered by the vast diversity of the gut microbiota and technical challenges in directly measuring DAMPA derived from bacterial metabolism. Although prior studies demonstrated that MTX can be depleted by gut bacteria or hydrolyzed *in vitro*, direct evidence linking specific gut bacteria to DAMPA production remains limited, as DAMPA is detected only at low levels.^23,25^ Consequently, the microbial contributors to DAMPA production remain poorly defined. Although microbial metabolites are increasingly recognized as key regulators of host physiology, the MTX metabolite DAMPA has long been regarded as an inert detoxification byproduct and therefore has not been considered a biologically active mediator in host–microbe–drug interactions.^26^ Thus, the physiological impact of DAMPA on host immune signaling and gut health has remained largely unexplored.

Here, we demonstrate that DAMPA is not an inert detoxification byproduct of MTX, rather it is a bioactive microbial metabolite that actively modulates intestinal epithelial and immune responses under inflammatory conditions. We identify gut bacterial species capable of producing DAMPA and show that DAMPA influences gut epithelial mitochondrial health and inflammatory responses both *in vitro* and *in vivo*. Our findings reveal an unrecognized functional role for microbial MTX metabolism in shaping host responses during intestinal inflammation and establish DAMPA as an effective mediator of host–microbe–drug interactions.

## RESULTS

### Microbial methotrexate metabolism to deoxyaminopteroic acid is reduced during intestinal inflammation

Methotrexate (MTX) is a folate mimic, structurally composed of a diaminopteridine ring linked to a p-aminobenzoyl group, and a glutamate tail (Figure 1A). Gut bacteria are linked to MTX metabolism, wherein the terminal glutamate moiety is cleaved to produce deoxyaminopteroic acid (DAMPA) via the action of the Carboxypeptidase G2 (CPDG2) enzyme (Figure 1A).^21,22^ CPDG2 is considered an exclusively bacterially produced enzyme, with no known mammalian counterpart that can deconjugate glutamate in MTX to produce DAMPA.^27^ Consistent with its microbial origin, CPDG2 was predominantly detected in the distal gastrointestinal tract, with strong signals observed in intestinal contents and only faint signals detected in intestinal tissue fractions in a western blot analysis (Figures S1A and S1B). Given the luminal localization of gut microbiota, we hypothesized that the weak CPDG2 signal in tissue fractions reflected residual luminal bacterial contamination rather than host expression.

**Figure 1.**
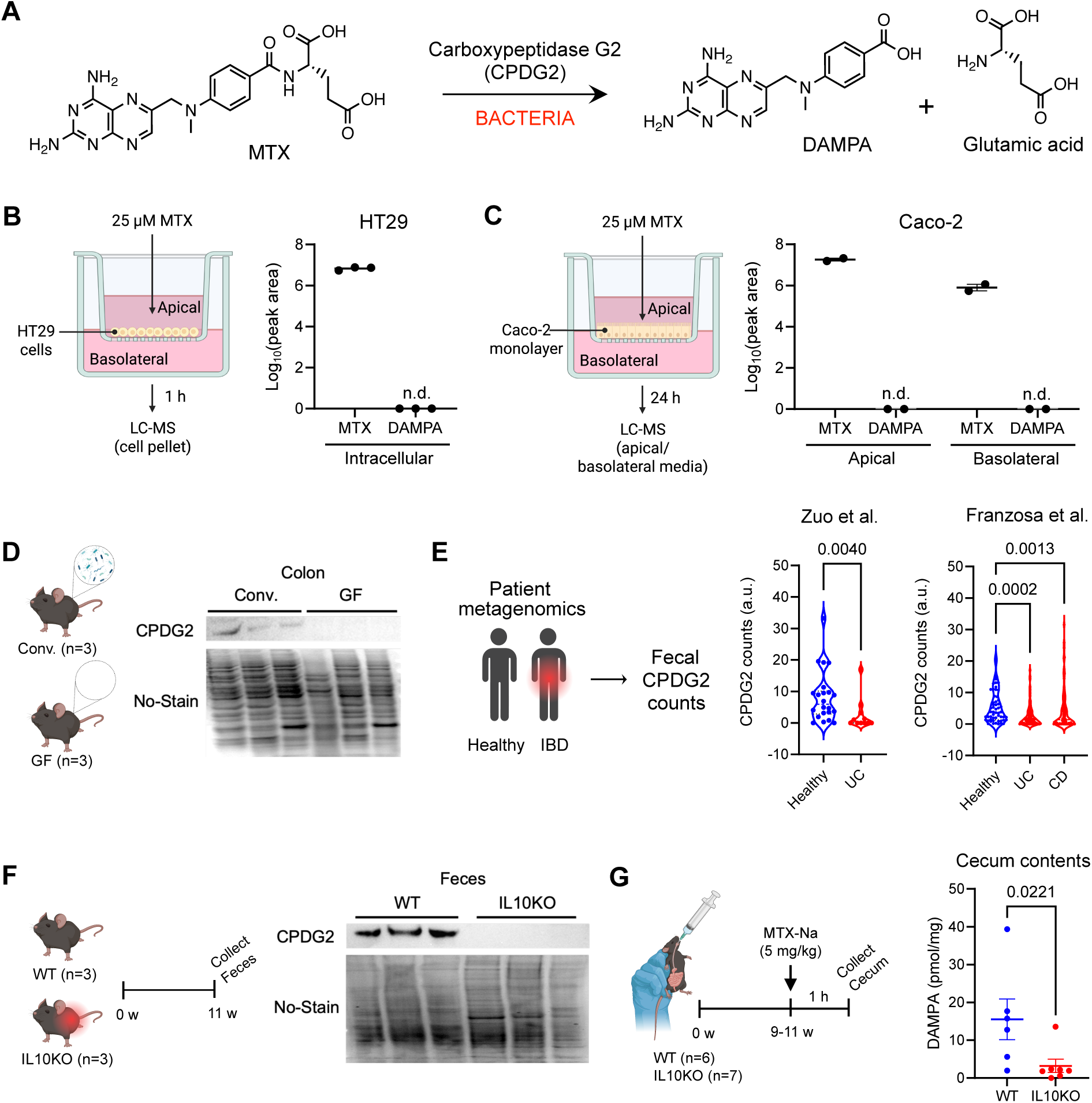
Microbiota-derived DAMPA production is attenuated by reduction of CPDG2 during gut inflammation. (A) Schematic of microbial MTX metabolism. (B) Apical MTX treatment in HT29 cells cultured on Transwell inserts. Cells were treated apically with MTX (25 µM) for 1 h, followed by DAMPA quantification in cell pellets by LC-MS (n.d. = not detected, n = 3). (C) Apical MTX treatment in differentiated Caco-2 monolayers cultured on Transwell inserts. Cells were treated apically with MTX (25 µM) for 24 h, and apical and basolateral medium were collected for DAMPA measurement by LC-MS (n.d. = not detected, n = 2). (D) CPDG2 immunoblot analysis of colon sections from 6–8-week-old conventional (Conv.) and germ-free (GF) mice (n = 3 per group). (E) CPDG2 gene abundance in fecal metagenomic datasets from healthy controls and patients with inflammatory bowel disease (IBD), including ulcerative colitis (UC) and Crohn’s disease (CD), normalized to total bacterial gene counts (Zuo et. al., healthy, n = 22; UC, n = 11, Mann-Whitney test; Franzosa et. al., healthy, n = 55; UC, n = 75; CD, n = 87, Kruskal-Wallis test). (F) Experimental scheme for fecal sample collection from WT and IL10KO mice at 11 weeks of age (n = 3 for each group). Immunoblot analysis of microbial CPDG2 protein in fecal samples from WT and IL10KO mice. Total protein loading was assessed using No-Stain total protein labeling. (n = 3 for each group). (G) Schematic of a single oral gavage of MTX-Na (5 mg/kg) administered to WT and IL10KO mice at 9-11 weeks of age, followed by cecum collection 1 h after administration. DAMPA levels in cecal contents measured by LC-MS (WT, n = 6; IL10KO, n = 7, Mann-Whitney test). All scatter dot plots are represented as mean ± SEM, with each data point representing an individual biological replicate. Violin plots show data distribution, with the central line indicating the median and dotted lines representing the interquartile range (25th–75th percentiles). Exact *p* values are indicated in each graph.

To directly test whether the host can contribute to DAMPA production, we exposed human intestinal epithelial cells in transwells to MTX and measured DAMPA levels (Figures 1B and 1C). Human colonic epithelial cells were incubated with 25 µM MTX, followed by extraction and quantification using a folate liquid chromatography/mass spectrometry (LC-MS) metabolomics method optimized for detection and quantification of folate and other polar metabolites in biological samples.^28^ This method allows for quantification of MTX and DAMPA with the limits of detection of 20 nM and 100 nM, respectively (Figures S2A and S2B). No detectable DAMPA was observed in HT29 human colonic epithelial cells in transwells following acute MTX exposure (Figure 1B). Similarly, DAMPA was not detected in differentiated human Caco-2 colonic epithelial monolayers after prolonged MTX treatment in either the apical or basolateral compartments (Figure 1C). Consistent with these *in vitro* findings, immunoblot analysis of colon segments with contents from conventional and germ-free animals showed CPDG2 detection in conventional mice with no detectable signal in germ-free mice (Figure 1D). Together, these results provide further evidence that DAMPA is produced by gut bacteria and not the host, establishing DAMPA as a microbiota-derived MTX metabolite in the large intestine.

Whether prevalent inflammation in the gut alters bacterial DAMPA production has not been previously examined. To determine whether intestinal inflammation affects microbial MTX metabolism, we assessed CPDG2 abundance and DAMPA production in human IBD patients and murine models of intestinal inflammation. Metagenomic analysis of human stool samples revealed a significant reduction in CPDG2 abundance in patients with IBD, including ulcerative colitis (UC) and Crohn’s disease (CD), compared to healthy controls (Figure 1E).^29,30^ To examine the impact of intestinal inflammation on microbial MTX metabolism *in vivo*, we utilized interleukin-10–deficient (IL10KO) mice, a well-established model of spontaneous colitis (Figure 1F).^31^ CPDG2 protein was detected in fecal samples from wild-type (WT) healthy controls but was undetectable in IL10KO mice (Figure 1F). To assess whether a reduction in CPDG2 enzyme abundance is indicative of low MTX metabolism, DAMPA production in the cecum was quantified in WT and IL10KO mice an hour after animals were administered MTX-Na (5 mg/kg) (Figure 1G). DAMPA concentrations were determined using standard curves generated in cecum contents to account for potential matrix effects during LC-MS analysis (Figures S2C and S2D). DAMPA production was markedly reduced in IL10KO mice by ∼80% as compared to the levels observed in WT controls (Figure 1G). Together, these results indicate that intestinal inflammation is associated with impaired microbial MTX metabolism and reduced DAMPA production in the lower gastrointestinal tract.

### *Clostridium asparagiforme*, a Firmicutes bacterium, contributes to microbial DAMPA production

To identify gut bacterial species capable of producing DAMPA, we first used a BLAST (Basic Local Alignment Search Tool)^32^ protein sequence analysis to identify CPDG2 homologs among 43 representative bacterial strains found in the human gut. Strains within the Firmicutes and Proteobacteria phyla exhibited higher CPDG2 alignment scores as compared to other bacteria (Figure 2A). Given that Firmicutes are abundant members of the human gut microbiota, whereas Proteobacteria are generally less abundant,^33^ we hypothesized that Firmicutes are major contributors to gut DAMPA production. Within Firmicutes, three clostridial species, *Clostridium symbiosum, Clostridium asparagiforme*, and *Clostridium scindens*, displayed high CPDG2 alignment scores. To validate our *in silico* results, we performed western blot analysis for CPDG2 protein in these species, and in representative bacteria from other major gut phyla including *Bifidobacterium longum* (Actinobacteria), *Bacteroides thetaiotaomicron* (Bacteroidetes), *Fusobacterium nucleatum* (Fusobacteria), *Escherichia coli Nissle 1917* (Proteobacteria), and *Akkermansia muciniphila* (Verrucomicrobia), as previously defined.^34^ Among the eight species tested, only *Clostridium symbiosum, Clostridium asparagiforme*, and *Clostridium scindens* exhibited CPDG2 expression, consistent with our *in silico* analyses, with *Clostridium asparagiforme (C. asparagiforme)* showing the strongest CPDG2 signal (Figure 2B). To determine whether CPDG2 expression correlated with DAMPA production, we cultured each strain anaerobically in the presence of MTX-Na (200 µM) and the CPDG2 cofactor ZnCl₂ (2 mM)^35^ in anaerobic medium for 1 h, and quantified DAMPA levels in bacterial pellets by LC-MS (Figure S3A). Of all species tested, *C. asparagiforme* showed a clear LC-MS peak for DAMPA, suggesting that *C. asparagiforme* can rapidly metabolize MTX to DAMPA (Figure 2C).

**Figure 2.**
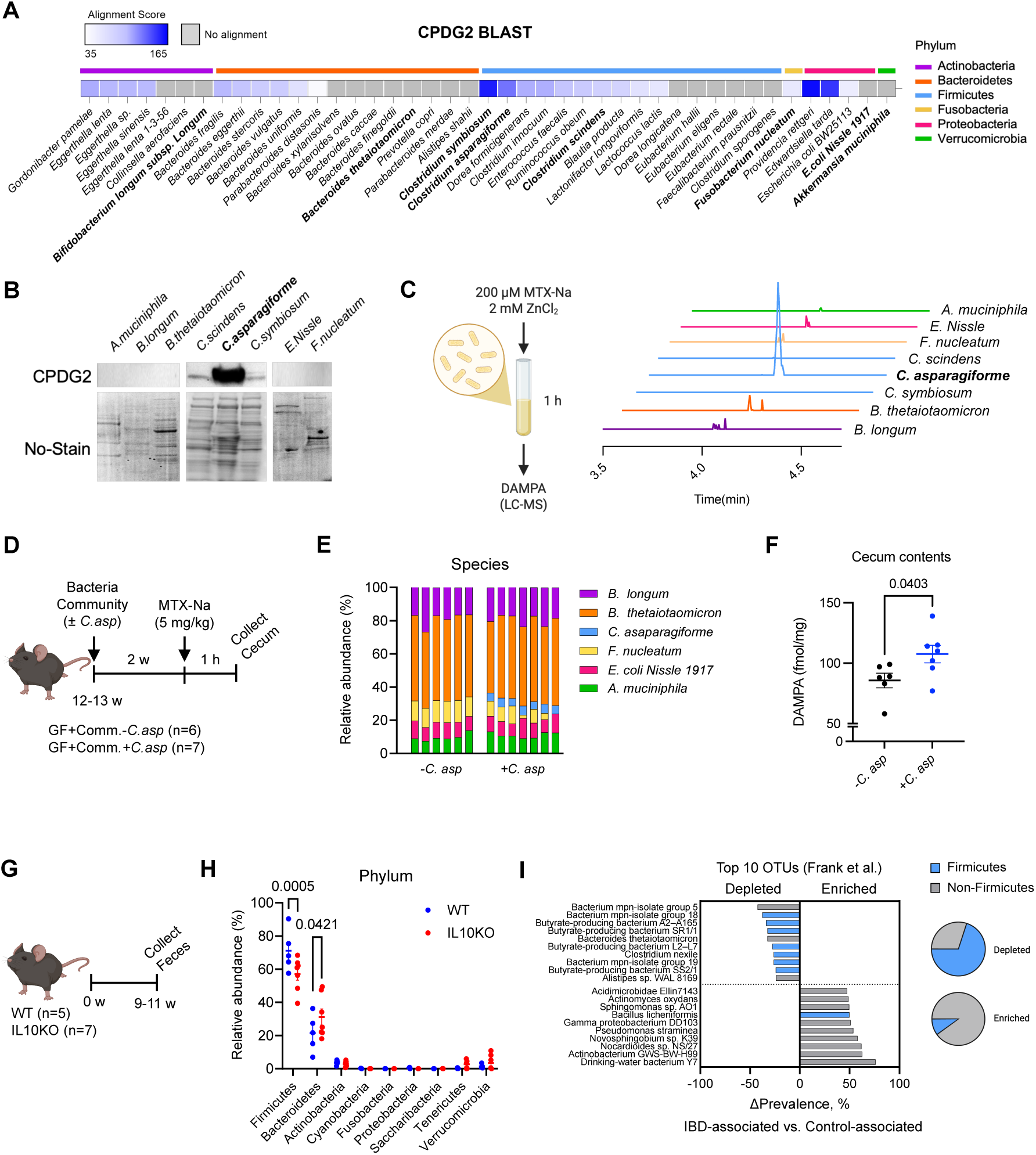
Clostridium asparagiforme produces DAMPA in vitro and in vivo. (A) Heatmap of CPDG2 BLAST alignment scores across 43 representative gut bacterial species spanning six phyla. Grey represents no alignment. (B) Immunoblot analysis of CPDG2 protein expression in cell pellets from eight bacterial species representing six gut bacterial phyla. Total protein loading was assessed using No-Stain total protein labeling. (C) Representative LC-MS chromatograms showing DAMPA in bacterial cell pellets following *in vitro* MTX exposure. Bacteria were cultured in Cullen–Haiser Gut (CHG) medium supplemented with MTX-Na (200 µM) and ZnCl₂ (2 mM) for 1 h in anaerobic chamber. (D) Schematic of the germ-free *in vivo* experiment. 12-13 weeks old germ-free mice were colonized with a defined bacterial consortium with or without *C. asparagiforme*, followed by oral administration of MTX-Na (5 mg/kg) and cecal content collection 1 h after gavage. (E) Relative abundance of cecal microbiota at the species level. Each bar represents biological replicates. (F) Cecal DAMPA levels measured by LC-MS after 1 h MTX-Na gavage (GF+Community (Comm.)-*C.asparagiforme (C. asp)*, n = 6; GF+Comm.+*C.asp*, n = 7, two-tailed Welch’s t test). (G) Experimental scheme for fecal sample collection from WT and IL10KO mice at 9-11 weeks of age. (H) Relative microbial composition at the phylum level in fecal samples from WT and IL10KO mice (WT, n = 5; IL10KO, n = 7, two-way ANOVA). (I) Top 10 Operational Taxonomic Units (OTUs) showing the largest differences in prevalence between IBD-associated and control-associated human microbiota profiles (adapted from Frank et al.). Phylum-level composition (%) of these OTUs is summarized in pie charts. All scatter dot plots are represented as mean ± SEM, with each data point representing an individual biological replicate. Exact *p* values are indicated in each graph.

To assess the *in vivo* ability of *C. asparagiforme* to produce DAMPA, germ-free mice were colonized with a defined bacterial consortium either lacking or including *C. asparagiforme* (Figure 2D). After microbiota stabilization, mice were administered a single oral dose of MTX-Na (5 mg/kg), and DAMPA levels were quantified in cecal contents one hour after gavage. Successful colonization was confirmed by 16S rRNA sequencing (Figure 2E). Despite comprising a minor fraction of the total microbiota composition (∼5.1%), presence of *C. asparagiforme* resulted in a significant ∼25% increase in DAMPA production compared to the control group (Figure 2F). Together, these results identify *C. asparagiforme* as a key contributor to microbial DAMPA production *in vivo*.

### DAMPA producers are decreased in abundance in gut inflammation

To determine whether inflammatory conditions alter microbiota potentially involved in MTX metabolism, we performed 16S rRNA sequencing in fecal samples from IL10KO mice and WT controls (Figure 2G). Principal coordinate analysis (PCoA) showed distinct clusters of microbiota composition in each group, with no significant differences in alpha diversity, suggesting differences in microbiota species composition and not richness between WT and IL10KO animals (Figures S4A and S4B). At the phylum level, the gut microbiota in both WT and IL10KO mice was dominated by Firmicutes and Bacteroidetes, consistent with previous reports (Figure 2H).^33^ Notably, IL10KO mice exhibited a significant reduction in the abundance of Firmicutes (∼14.0%) with a corresponding increase in Bacteroidetes (∼9.6%), indicating inflammation-associated restructuring of the gut microbiome that would decrease DAMPA production. This is consistent with previous studies reporting a reduction in the relative abundances of Firmicutes in human IBD cohorts.^36,37^ Analysis of the PRISM and HMP2 cohorts show a consensus decrease in Firmicutes in IBD patient stool samples compared to healthy controls (Table S1).^38^ When comparing microbiota profiles from IBD patients to those from healthy controls, 7 of the 10 bacterial taxa showing the largest decreases in prevalence were classified as Firmicutes. (Figure 2I; adapted from Frank et al.^36^).

While Bacteroidetes-associated taxa were broadly increased in IL10KO mice, Firmicutes-associated classes and families showed variable, bidirectional shifts, suggesting non-uniform restructuring within this phylum (Figures S4C and S4D). Prior human studies have shown a selective reduction in *Clostridia*, a major class within Firmicutes phylum, in patients with Crohn’s disease.^39,40^ A recent study identified a decrease in *C. asparagiforme* in IBD patients.^41^ Together, these results suggest that inflammation-associated changes in Firmicutes, specifically a decrease in *Clostridia and C. asparagiforme* may contribute to impaired DAMPA production in the gut.

### DAMPA and MTX exert dichotomous modulation of mitochondria in intestinal epithelial cells under inflammatory conditions

Next, we wanted to investigate whether a decrease in bacterial MTX to DAMPA production in the context of gut inflammation has any physiological effects. To address this, we utilized HT29 human colonic epithelial cells treated with TNFα for 24 h to mimic inflammatory conditions *in vitro* (Figure 3A).^42^ We then treated cells with DAMPA or MTX in the continued presence of TNFα overnight, followed by untargeted proteomic analyses of cells harvested the next day. Principal component analysis (PCA) revealed a clear separation of the DMSO-treated controls and drug-treated cells, indicating substantial proteomic remodeling upon MTX or DAMPA exposure (Figure 3B). DAMPA treatment resulted in widespread changes in protein abundance compared to DMSO controls, as visualized in the volcano plot (Figure 3C). In a DAMPA and MTX comparison, among the significantly upregulated proteins, Dihydrofolate Reductase (DHFR) abundance was prominently increased in the DAMPA treatment group as compared to MTX, consistent with the inhibitory effect of MTX on DHFR (Figure 3D).^43^ Gene ontology (GO) enrichment analysis^44^ of proteins upregulated by DAMPA revealed significant enrichment of biological processes associated with cellular stress adaptation and immune-related responses (Figure 3E). In parallel, comparison of DAMPA versus MTX treatment highlighted differential regulation of pathways linked to cellular responses to immune signaling (Figure 3F). Cellular component GO analysis demonstrated a significant enrichment of mitochondria-associated proteins following DAMPA treatment, overall indicating a selective impact of this microbial MTX metabolite on stress adaptation and mitochondrial health (Figure 3G). Consistent with proteomic findings, DAMPA increased expression of the Ubiquinol-Cytochrome C Oxidoreductase Binding Protein (UQCRB), a component of mitochondrial complex III, at the mRNA level compared to DMSO controls (Figure S5A), supporting selective remodeling of the electron transport chain.

**Figure 3.**
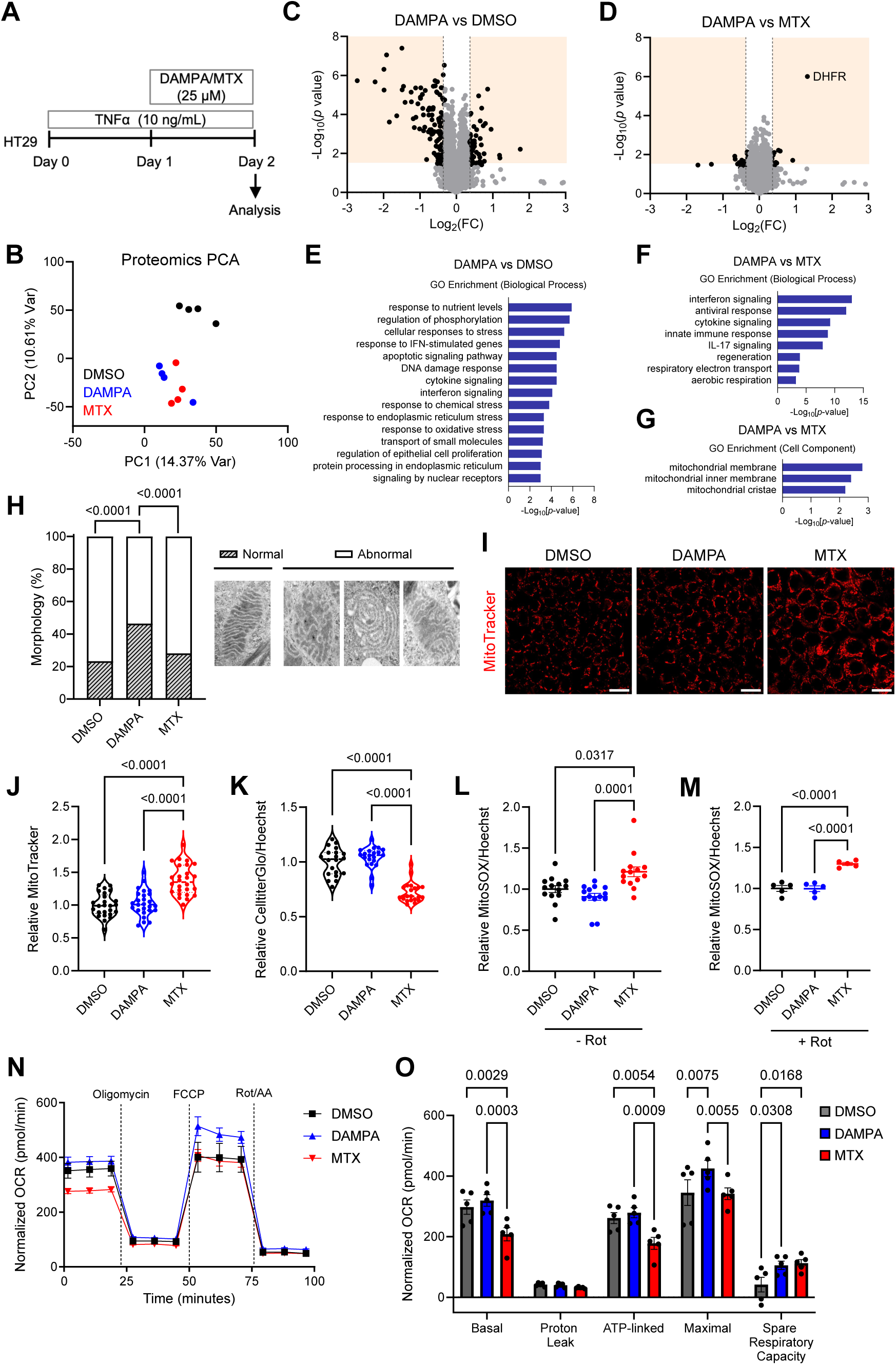
Dichotomous roles of DAMPA and MTX on mitochondrial health during intestinal inflammation. (A) Schematic of HT29 cell experimental design. Cells were pretreated with TNFα (10 ng/mL) for 24 h, followed by DAMPA (25 µM) or MTX (25 µM) treatment for an additional 24 h. (B) Principal component analysis (PCA) plot of proteomic profiles from HT29 cells (n = 4 per group). A total of 8,036 proteins were analyzed. (C and D) Volcano plots showing differentially abundant proteins. The –Log_10_(*p* value) is plotted against Log_2_(fold change). Differentially abundant proteins (fold change > 1.25) are indicated by black dots (dotted lines denote cutoffs). Comparisons shown are DAMPA vs DMSO (C) and DAMPA vs MTX (D). (E-G) Gene Ontology (GO) enrichment analyses of DAMPA-upregulated proteins (fold change > 1.25, *p* < 0.05). GO biological process enrichment for DAMPA vs DMSO (E), GO biological process enrichment for DAMPA vs MTX (F), and GO cellular component enrichment for DAMPA vs MTX (G). (H) Proportion of mitochondrial morphologies assessed by transmission electron microscopy (TEM). Mitochondria were classified as normal or abnormal based on cristae morphology. (DMSO, n = 30; DAMPA, n = 28; MTX, n = 32; n represents individual mitochondria, chi-square test). Representative TEM images of normal and abnormal cristae morphologies are shown. (I and J) MitoTracker staining. (I) Representative MitoTracker-stained images of HT29 cells treated with DMSO, DAMPA, or MTX. Scale bar, 20 µm. (J) Relative MitoTracker fluorescence intensity quantified by plate reader. MitoTracker signal (Ex 579 nm, Em 599 nm) was normalized to nuclear staining (DAPI or Hoechst 33342) (DMSO, n = 25; DAMPA, n = 26; MTX, n = 26, one-way ANOVA). (K) Cellular ATP levels measured by CellTiter-Glo. Luminescence was measured by plate reader and normalized to Hoechst 33342 fluorescence (DMSO, n = 20; DAMPA, n = 21; MTX, n = 21, one-way ANOVA). (L and M) Relative mitochondrial superoxide levels assessed by MitoSOX fluorescence without (L) or with (M) rotenone (Rot., 250 nM). Fluorescence intensity was quantified using ImageJ and normalized to Hoechst 33342. (L) n = 14, Kruskal–Wallis test; (M) n = 5, one-way ANOVA. (N and O) Oxygen consumption rate (OCR) measured by Agilent Seahorse XFe24 analyzer. OCR values were normalized to mitochondrial content as determined by MitoTracker staining (n = 5 per group, two-way ANOVA). All scatter dot plots and bar graphs are represented as mean ± SEM, with each data point representing an individual biological replicate. Violin plots depict data distribution, with the central line indicating the median and dotted lines representing the interquartile range (25th–75th percentiles). Exact *p* values are shown in the graphs.

Based on the proteomics results, we next examined whether DAMPA affects mitochondrial morphology under inflammatory conditions. We performed transmission electron microscopy (TEM) on HT29 cells treated with DAMPA or MTX under inflammatory stimuli (Figure 3A). Both DMSO– and MTX-treated cells exhibited substantial disruption of mitochondrial cristae architecture, with ∼80% of mitochondria displaying abnormal cristae morphology, including onion-like and honeycomb-like structures, and cristae loss (Figure 3H). These cristae abnormalities are characteristic of dysfunctional or defective mitochondria across multiple tissues and organisms.^45–47^ In contrast, DAMPA-treated cells showed a significantly higher proportion of mitochondria with preserved lamellar cristae architecture, indicative of normal mitochondrial morphology. These results suggest that DAMPA, but not MTX, can improve mitochondrial morphology derangements induced by inflammation in gut epithelial cells.

To further assess mitochondrial health metrics, we examined mitochondrial membrane potential and network using MitoTracker.^42^ MTX-treated cells exhibited increased MitoTracker fluorescence intensity compared to both DAMPA– and DMSO-treated cells, suggesting altered mitochondrial membrane potential and increased mitochondrial content under inflammatory stress (Figures 3I and 3J). Combined with our previous results (Figure 3H), this suggests accumulation of dysfunctional mitochondria in MTX-treated cells compared to DAMPA. Mitochondrial Network Analysis (MiNA), together with qPCR analyses for key mitochondrial biogenesis, fusion, and fission markers revealed that MTX increased mitochondrial biogenesis and fusion, while DAMPA reduced Dynamin-1-like (*DNM1L)* expression, a key regulator of mitochondrial fission (Figures S5B-S5I).^48^

To determine whether these structural and dynamic changes affect mitochondrial function, we measured cellular ATP and mitochondrial superoxide levels. MTX treatment significantly reduced ATP production by ∼30% and markedly increased mitochondrial superoxide levels, in the presence or absence of rotenone, a complex I inhibitor that induces mitochondrial ROS generation (Figures 3K-3M and S6).^49^ While MTX had deleterious effects on mitochondrial structure and health metrics, DAMPA did not show any marked improvement when compared to DMSO control. Therefore, we next measured mitochondrial oxidative phosphorylation capacity using Seahorse analysis to measure oxygen consumption rate (OCR). MTX significantly impaired basal and ATP-linked mitochondrial respiration compared to DAMPA and DMSO controls (Figures 3N and 3O). Strikingly, DAMPA treatment significantly increased maximal OCR when compared to MTX and DMSO control and enhanced spare respiratory capacity, indicating improved mitochondrial functional adaptability under inflammatory stress (Figures 3N and 3O). Overall, these findings indicate that while MTX compromises mitochondrial integrity and function in intestinal epithelial cells, its microbiota-derived metabolite DAMPA improves mitochondrial structure and enhances functional resilience toward oxidative phosphorylation under inflammatory conditions.

### DAMPA induces autophagy in intestinal epithelial cells under inflammatory stress

Our TEM analyses revealed a significant accumulation of autophagosome-like structures in intestinal epithelial cells treated with DAMPA (Figures 4A and 4B). During prolonged cellular stress conditions such as inflammation, endoplasmic reticulum (ER) stress can activate autophagy as a protective mechanism to clear damaged proteins and dysfunctional organelles.^50–52^ Consistently, our proteomics results supported that DAMPA significantly enriched pathways associated with the ER, including ER stress response and protein processing (Figure 3E). To validate these observations at the molecular level, we assessed lipidated Microtubule-Associated Protein 1 Light Chain 3 Beta (LC3B-II), a standard marker of autophagosomes,^53,54^ by western blot analysis. DAMPA treatment showed a significant increase in LC3B-II levels compared with both MTX and DMSO controls (Figure 4C), suggesting that DAMPA promotes autophagy under inflammatory conditions. Autophagy requires lysosomal fusion to enable clearance of damaged proteins and organelles.^55^ Given that DAMPA preserved mitochondrial morphology and enhanced respiratory capacity (Figures 3H, 3N and 3O), we next asked whether DAMPA-induced autophagy is associated with mitochondrial quality control. To this end, we co-stained lysosomes and mitochondria using LysoTracker and MitoTracker in TNFα-stimulated HT29 cells treated with DAMPA or MTX (Figure 3A). Quantification of mitochondria-lysosome tethering revealed a significant increase in mitochondria-lysosome interactions in DAMPA-treated cells (Figures 4D and 4E).^56,57^ This increase occurred without changes in lysosome abundance or functional markers, as indicated by comparable LysoTracker intensity and levels of Lysosome-Associated Membrane Protein 1 (LAMP1) and ATPase H+ transporting V1 subunit A (ATP6V1A) expression across treatment groups (Figures S7A–S7C).^58,59^

**Figure 4.**
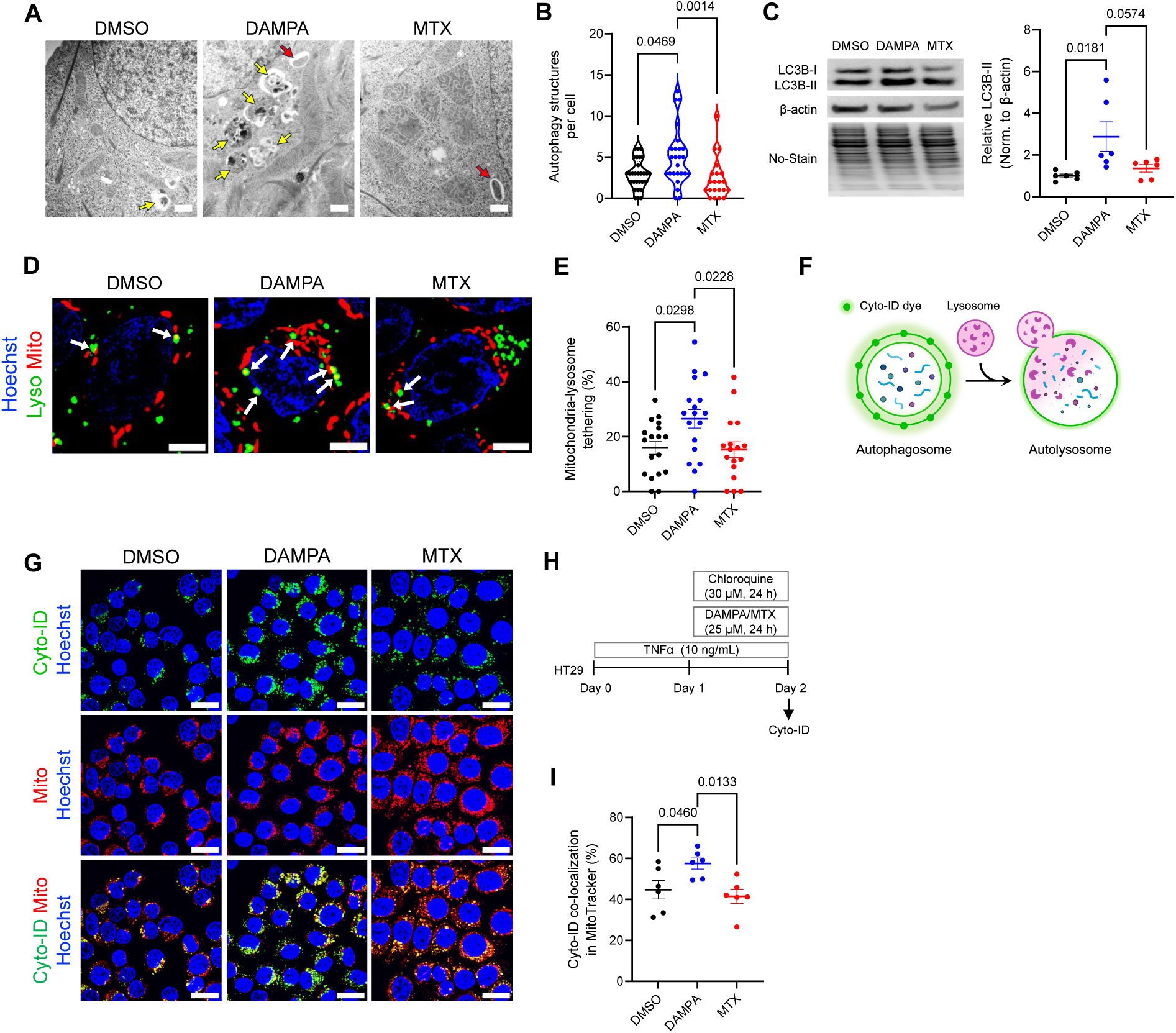
DAMPA promotes autophagy in intestinal epithelial cells under inflammatory stress. (A) Representative transmission electron microscopy (TEM) images of HT29 cells treated with DAMPA or MTX under TNFα-induced inflammatory conditions. Arrows indicate autophagy structures (red, autophagosomes; yellow, autolysosomes). Scale bar, 600 nm. (B) Quantification of autophagic structures (phagophores, autophagosomes, and autolysosomes) per cell. (DMSO, n = 29; DAMPA, n = 25; MTX, n = 21, Kruskal-Wallis test) (C) LC3B-II protein abundance in TNFα-treated HT29 cells exposed to DAMPA or MTX. LC3B-II levels were normalized to β-actin. (n = 6 per group, two-tailed Welch’s t test) (D and E) Co-staining of mitochondria (MitoTracker) and lysosomes (LysoTracker) in HT29 cells. (D) Representative images of MitoTracker and LysoTracker co-staining. Arrows indicate mitochondria-lysosome contacts. Nuclei were stained with Hoechst 33342. Scale bar, 5 µm. (E) Quantification of mitochondria-lysosome contacts per lysosome per cell. (DMSO, n = 18; DAMPA, n = 18; MTX, n = 17, one-way ANOVA) (F-I) Autophagic compartment and mitochondrial staining in HT29 cells using Cyto-ID (green) and MitoTracker (red). (F) Schematic of Cyto-ID staining. (G) Representative images showing autophagic compartments (Cyto-ID, green), mitochondria (MitoTracker, red), and nuclei (Hoechst 33342, blue). Scale bar, 20 µm. (H) Experimental workflow for Cyto-ID staining in HT29 cells. Cells were treated with MTX (25 µM) or DAMPA (25 µM) along with chloroquine (30 µM) to inhibit lysosomal activity under inflammatory conditions induced by TNFα (10 ng/mL). (I) Quantification of Cyto-ID and MitoTracker colocalization. The overlapping area was normalized to total mitochondrial area and expressed as the percentage of mitochondrial area colocalized with Cyto-ID (n = 6 per group, one-way ANOVA). All scatter dot plots are represented as mean ± SEM, with each data point representing an individual biological replicate. The violin plot depicts data distribution, with the central line indicating the median and dotted lines representing the interquartile range (25th–75th percentiles). Exact *p* values are shown in the graphs.

To validate whether increased lysosome-mitochondria tethering reflects mitophagy, we quantified autophagic mitochondria by utilizing Cyto-ID staining to label autophagic organelles including autophagosomes and autolysosomes (Figures 4F-4I).^60^ Cells were treated with TNFα then co-treated with chloroquine and DAMPA or MTX to measure autophagy flux (Figure 4H).^61^ Quantification of mitochondria and Cyto-ID colocalization revealed a significant increase in Cyto-ID positive mitochondria in DAMPA-treated cells, suggesting that DAMPA induces mitophagy in intestinal epithelial cells (Figures 4G and 4I). We next assessed PTEN-Induced Kinase 1 (PINK1), a marker of Parkin-dependent mitophagy.^62,63^ We did not observe detectable changes in PINK1 protein abundance by western blot or immunostaining analyses, suggesting limited involvement of the canonical Parkin-dependent mitophagy pathway (Figures S7D and S7E). Overall, these results suggest that DAMPA induces autophagy as a broader cytoprotective stress response linked to ER stress and potentially mitochondrial clearance through a Parkin-independent pathway, without an increase in lysosomal degradative capacity.

### DAMPA modulates STAT3 signaling through folate receptor binding

Next, we sought to investigate the mechanism by which DAMPA improves mitochondrial health and promotes autophagy during gut inflammation. We first investigated whether DAMPA is imported into gut epithelial cells to interact with intracellular proteins or organelles directly. In our *in vitro* experiments, DAMPA was not detected within gut epithelial cells following apical treatment in HT29 cells, nor was it detected in the basolateral compartment of differentiated Caco-2 monolayers after apical DAMPA exposure (Figures S8B and S8C). Consistently, upon oral gavage of MTX in mice (Figure 1J), DAMPA was detected exclusively in intestinal luminal contents, but not in serum (data not shown). These observations suggest that DAMPA is not transported into epithelial cells but instead may act as an extracellular signaling molecule when generated in the intestinal lumen by microbial MTX metabolism.

Given the structural similarity between DAMPA and folic acid (Figure S8A), we hypothesized that DAMPA may interact with folate-associated receptors or transporters. Mammalian cells utilize three major membrane-bound folate interacting proteins: the Reduced Folate Carrier (RFC), the Proton-Coupled Folate Transporter (PCFT), and the Folate Receptor (FR).^64^ Among these, FR is unique in its ability to initiate intracellular signaling cascades upon ligand binding.^64^ Mammals have 4 FR isoforms, FRα, FRβ, FRγ, and FRδ. Considering that FRα showing highest expression in the gastrointestinal tract,^65,66^ we hypothesized that DAMPA can bind and activate FRα on gut epithelial cells. Structural studies have shown that the FRα recognizes folate primarily through its pteroate moiety, whereas the glutamate tail remains solvent-exposed and does not substantially contribute to receptor affinity.^67^ Because DAMPA preserves the pteroate core structure of folate (Figure S8A), its ability to bind FRα is structurally plausible. Consistent with this model, molecular docking analysis^68^ using the human FRα predicted that DAMPA binds tightly within the FRα binding pocket, with an affinity greater than that of MTX and folic acid (Figure 5A). To test whether DAMPA binds FRα, we performed a competitive binding assay using isolated cell membrane fractions from HT29 cells. Briefly, membranes were incubated with labeled folic acid (folic acid-d2), and its displacement by DAMPA was measured via LC-MS. This assay showed that DAMPA binds FRα on gut epithelial cells with high affinity (EC_50_ = 2.07 nM) (Figure 5B), supporting FRα-mediated signaling as a key mechanism for DAMPA in intestinal epithelial cells.

**Figure 5.**
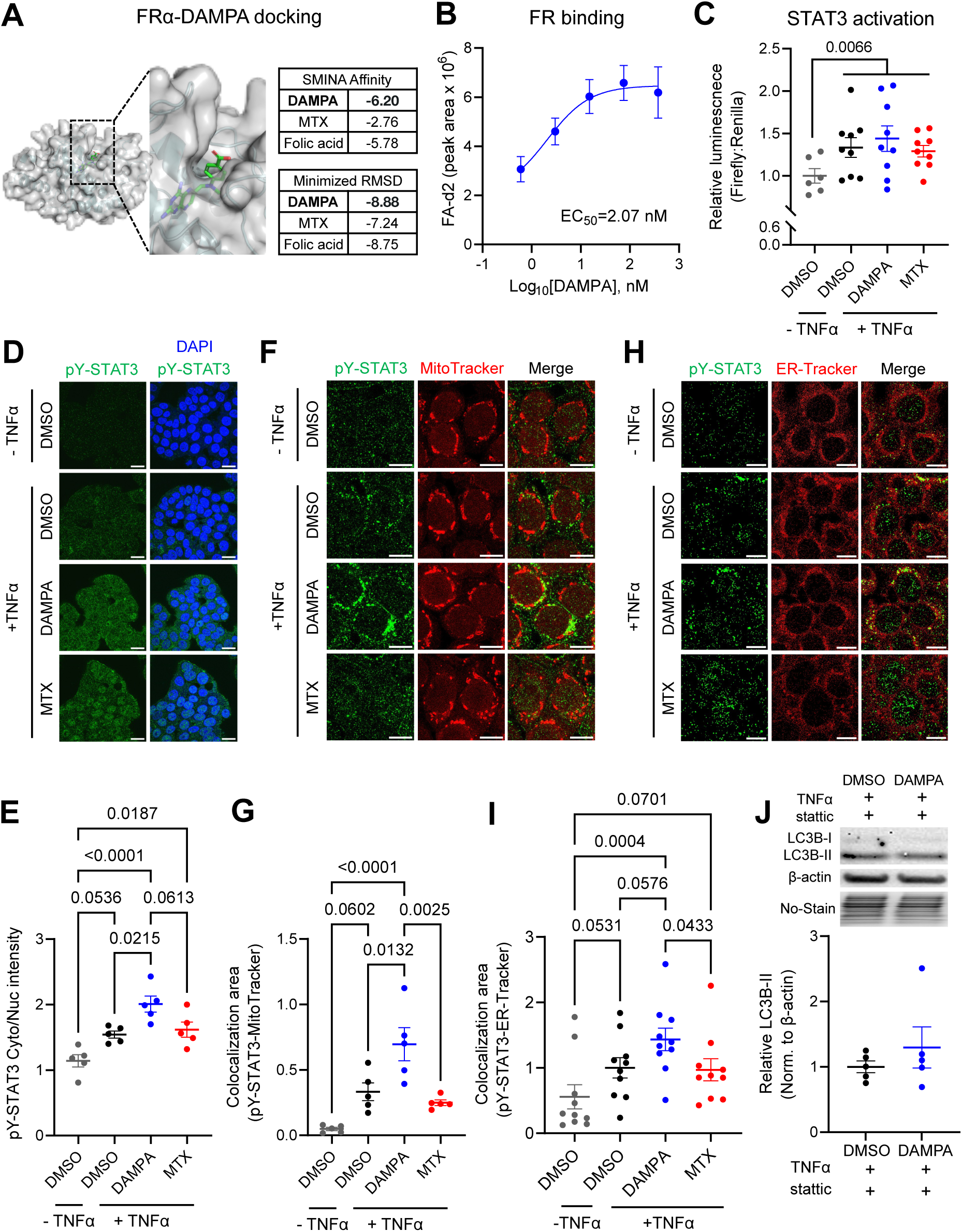
DAMPA binds folate receptor to promote cytoplasmic STAT3 localization. (A) *In silico* docking of DAMPA with human Folate Receptor Alpha (FRα; PDB: 4LRH) using DiffDock-L. Lower SMINA affinity indicates stronger predicted binding. Lower Minimized RMSD minimized affinity indicates more stable and energetically favorable binding. (B) Competitive binding assay of DAMPA to folate receptor using membrane fractions isolated from HT29 cells. Folic acid-d2 (15 nM) was used as a competitor. Binding curves were generated by nonlinear regression of peak area under the curve (AUC) measured by LC-MS (n = 5 per group). (C) STAT3 activation measured by a dual-luciferase reporter assay in HT29 cells treated with DAMPA or MTX treatment in the absence and presence of TNFα (DMSO without TNFα; n = 6, all other groups; n = 9 per group, two-way ANOVA). (D and E) Phosphorylated STAT3 at Tyr705 (pY-STAT3) staining in HT29 cells treated with DAMPA or MTX in the presence or absence of TNFα. (D) Representative images. Scale bar, 20 µm. (E) Ratio of cytoplasmic to nuclear pY-STAT3 fluorescence intensity quantified by ImageJ (n = 5 per group, two-way ANOVA). (F and G) Co-staining of pY-STAT3 and mitochondria in HT29 cells. (F) Representative images showing pY-STAT3 and MitoTracker colocalization. Scale bar, 10 µm. (G) Quantification of pY-STAT3–mitochondria colocalization area by Image J. Colocalization area was normalized to cell number (DAPI) (n = 5 per group, two-way ANOVA). (H and I) Co-staining of pY-STAT3 and the endoplasmic reticulum (ER) in HT29 cells. (H) Representative images showing pY-STAT3 and ER-Tracker colocalization. Scale bar, 10 µm. (I) Colocalization area was quantified using Image J and normalized to total cell number based on DAPI staining (n = 10 per group, two-way ANOVA). (J) LC3B-II expression in HT29 cells treated with the STAT3 inhibitor stattic (5 µM), 1 h prior to DAMPA exposure under TNFα stimulation. LC3B-II levels were normalized to β-actin (n = 5 per group). All scatter dot plots are represented as mean ± SEM, with each data point representing an individual biological replicate. Exact *p* values are indicated in each graph.

The Signal Transducer and Activator of Transcription 3 (STAT3) is a well-characterized downstream signaling effector engaged by FRα signaling.^69^ Folic acid and folinic acid can activate STAT3 through FRα in a Janus Kinase (JAK)-dependent manner.^69^ STAT3 signaling is highly context dependent.^70–73^ In addition to its canonical role in regulating inflammatory gene expression, STAT3 has been reported to localize to cytoplasmic compartments, including mitochondria and the ER, where it can modulate mitochondrial respiration and stress adaptation.^72,73^ Given the complex and context-dependent roles of STAT3 in inflammation, we first sought to define STAT3 activation under TNFα exposure in HT29 intestinal epithelial cells using a STAT3 luciferase-based reporter assay. We observed that TNFα treatment increased STAT3 activation across all groups compared to DMSO (Figure 5C). Because STAT3 function is critically influenced by its subcellular localization, we next investigated whether DAMPA modulates STAT3 localization under TNFα-induced inflammatory conditions. STAT3 contains two major phosphorylation sites, Tyr705 and Ser727.^74^ Tyr705 is canonically associated with STAT3 dimerization and transcriptional activity, whereas Ser727 has been linked to mitochondrial functions, including regulation of the electron transport chain and reactive oxygen species production.^75,76^ We performed immunocytochemical analysis of Ser727 and Tyr705 phosphorylated STAT3 (p-STAT3), both considered the activated form of STAT3. Notably, DAMPA treatment selectively increased cytoplasmic Tyr705 p-STAT3 (pY-STAT3) localization compared to MTX and DMSO controls, whereas Ser727 phosphorylation (pS-STAT3) was largely restricted to the nucleus (Figures 5D and 5E, S9). This suggests that DAMPA modulates STAT3 subcytoplasmic localization rather than nuclear localization linked to transcriptional activation.

p– STAT3 has been reported to localize to cytoplasmic compartments, including mitochondria and the ER, where it can regulate mitochondrial function and stress responses.^72,73^ At these sites, p-STAT3 supports mitochondrial respiratory function and coordinates ER stress responses, contributing to mitochondrial integrity.^77^ To determine whether DAMPA alters pY-STAT3 distribution within these cytoplasmic compartments, we co-stained pY-STAT3 with mitochondria using MitoTracker or with the ER using ER-Tracker. DAMPA treatment significantly increased the colocalization of pY-STAT3 with both mitochondria and the ER, compared to control (Figures 5F-5I). To further examine whether STAT3 signaling contributes to DAMPA-induced autophagy, we treated HT29 cells with the STAT3 inhibitor, stattic,^78^ with concomitant exposure to DAMPA. Stattic significantly abolished the DAMPA-mediated increase in LC3B-II protein levels (Figure 5J), indicating that STAT3 activity contributes to autophagy induction by DAMPA. Collectively, these results suggest that DAMPA engages FR–STAT3 signaling and promotes STAT3 subcellular localization to the mitochondria and the ER, which may support mitochondrial stress adaptation through autophagy under inflammatory conditions.

### Oral DAMPA administration attenuates inflammation *in vivo*

In IBD, intestinal epithelial cells exhibit impaired mitochondrial integrity, characterized by altered expression of mitochondria-related genes, disrupted cristae morphology, reduced oxidative metabolism, and increased ROS, collectively linked to the initiation and progression of gut inflammation.^79,80^ Proteomic analyses in HT29 cells revealed that DAMPA modulated pathways associated with inflammatory responses (Figures 3E and 3F). Together with our observations that DAMPA preserves mitochondrial morphology and function under inflammatory conditions, this suggests that DAMPA may exert anti-inflammatory effects by supporting mitochondrial stress adaptation in intestinal epithelial cells.

Based on these findings, we next tested whether DAMPA administration could ameliorate intestinal inflammation *in vivo* using IL10KO mice. Given that DAMPA remains in the intestinal lumen and is not transported into epithelial cells, IL10KO mice were administered DAMPA by oral gavage (4 mg/kg) daily for 50 days (Figure 6A). DAMPA treatment did not alter body weight, colon length, and spleen weight compared to vehicle controls (Figure S10). However, histological analysis of colonic tissues revealed a significant increase in crypt length in DAMPA-treated mice, indicating improved epithelial integrity (Figures 6B and 6C). DAMPA-treated mice also showed increased goblet cell abundance, further supporting enhanced mucosal homeostasis in the colon (Figure 6D). In parallel, neutrophil infiltration in the colon was significantly reduced following DAMPA administration (Figure 6E). Consistent with these histological findings, fecal lipocalin-2 (LCN-2), a sensitive marker of intestinal inflammation,^81^ was reduced by ∼60% in DAMPA-treated mice at day 35 compared with DMSO-treated controls (Figure 6F). While LCN-2 levels continued to rise significantly over time in DMSO-treated mice, DAMPA treatment blunted this increase, maintaining significantly lower LCN-2 levels at day 50 (Figure 6F). Serum cytokine analysis further demonstrated >50% reduction in levels of pro-inflammatory cytokines, including Granulocyte-Macrophage Colony-Stimulating Factor (GM-CSF), Interleukin-6 (IL-6), and Interferon-Alpha (IFN-α),^82^ in the DAMPA treatment group compared to control (Figures 6G). Proteomic analysis of colon tissues revealed that FUN14 Domain Containing 1 (FUNDC1), a protein involved in Parkin-independent mitophagy pathway,^83^ was significantly upregulated by DAMPA (Figure 6H). Western blot analysis validated the increased in FUNDC1 protein levels in colon tissues from DAMPA-treated mice (Figure 6I). To determine whether these anti-inflammatory effects were associated with gut microbial composition changes, we performed 16S rRNA sequencing of fecal samples. No significant changes were observed in overall microbiota composition or alpha diversity (Figures 6J-6M), indicating that the anti-inflammatory effects of DAMPA occur independently of gut microbiome shifts. Together, these results demonstrate that DAMPA induces autophagy in the gut and attenuates intestinal inflammation *in vivo* independently of microbiome remodeling (Figure 6N).

**Figure 6.**
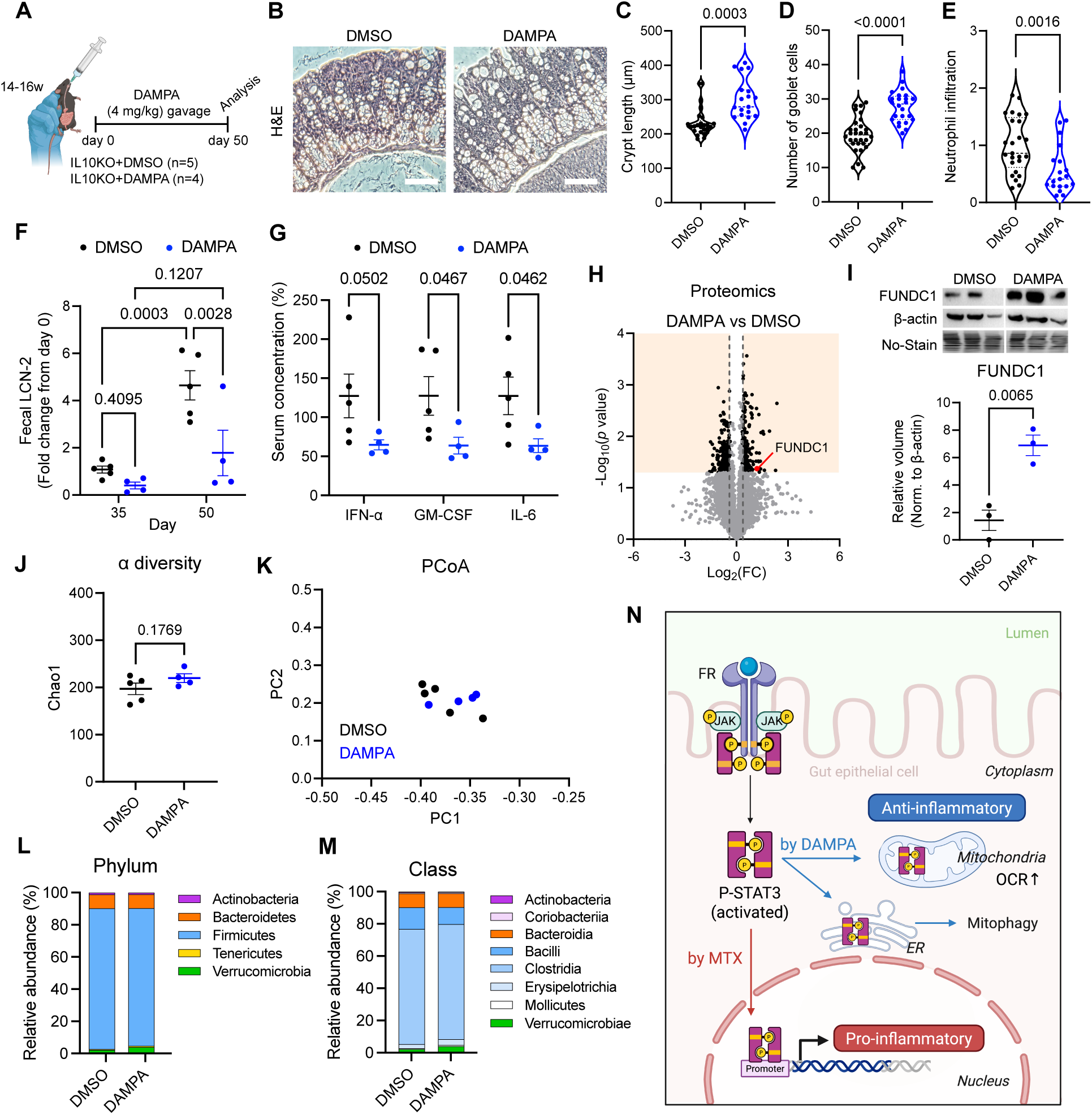
DAMPA administration attenuates intestinal inflammation in IL10KO mice. (A) Schematic of a daily oral gavage of DAMPA (4 mg/kg) administered to IL10KO mice (14-16 weeks old) for 50 days. (B-E) Histological analysis of colonic tissues at day 50. (B) Representative hematoxylin and eosin (H&E)-stained colon sections. Scale bar, 200 µm. (C) Crypt length quantification (DMSO, n = 25; DAMPA, n = 19, two-tailed Welch’s t test), (D) the number of goblet cells per crypt (DMSO, n = 30; DAMPA, n = 24, two-tailed Welch’s t test), and (E) neutrophil infiltration normalized to crypt area and expressed relative to the DMSO control (DMSO, n = 25; DAMPA, n = 19, Mann-Whitney test). Each n represents one crypt. 4-6 crypts per mouse were analyzed. (F) Fecal lipocalin-2 (LCN-2) levels measured by ELISA in samples collected at day 35 and day 50. Each n represents one mouse (DMSO, n = 5; DAMPA, n = 4, two-way ANOVA). (G) Serum pro-inflammatory cytokine (GM-CSF, IFN-α, and IL-6) levels measured by multiplex bead-based immunoassay. Each n represents one mouse (DMSO, n = 5; DAMPA, n = 4, two-way ANOVA). (H) Volcano plot showing differentially expressed proteins (DEPs) from the coon tissue proteomics analysis. The colored background indicates proteins with *p* < 0.05. (I) Western blot analysis of FUNDC1 protein levels in colon tissues (DMSO, n = 3; DAMPA, n = 3, two-tailed Welch’s t test). (J-M) 16S rRNA sequencing of fecal samples collected at day 50 (DMSO, n = 5; DAMPA, n = 4). (J) Alpha diversity (Chao1 index). (K) Principal coordinate analysis (PCoA). (L and M) Average relative abundance of microbial composition at (L) the phylum and (M) the class levels. (N) Proposed model of DAMPA-induced FR–STAT3 signaling. DAMPA binding to folate receptor promotes STAT3 localization to cytoplasmic compartments, including mitochondria and the endoplasmic reticulum, which may support mitochondrial function and mitophagy under inflammatory conditions. All scatter dot plots are represented as mean ± SEM. Violin plots depict the distribution of the data. The central line indicates the median, and dotted lines indicate the interquartile range (25th–75th percentiles). Each data point represents one crypt (C-E) or individual biological replicate (F, G, and I-K). Exact *p* values are indicated in each graph.

### DAMPA counteracts mitochondrial pathways disrupted in human IBD

To investigate the molecular pathways underlying the anti-inflammatory effects of DAMPA *in vivo*, we performed untargeted proteomic analysis on colon tissues from IL10KO mice treated with DAMPA or DMSO control for 50 days (Figure 7A). Among 412 differentially expressed proteins, 66 were annotated to the Gene Ontology (GO) term “mitochondrion”. GO biological process enrichment analysis of these 66 proteins revealed a significant enrichment in mitochondrial metabolic pathways, with the electron transport chain emerging as the most significantly enriched term, followed by lipid metabolism, the tricarboxylic acid cycle (TCA), and the urea cycle (Figure 7B).

**Figure 7.**
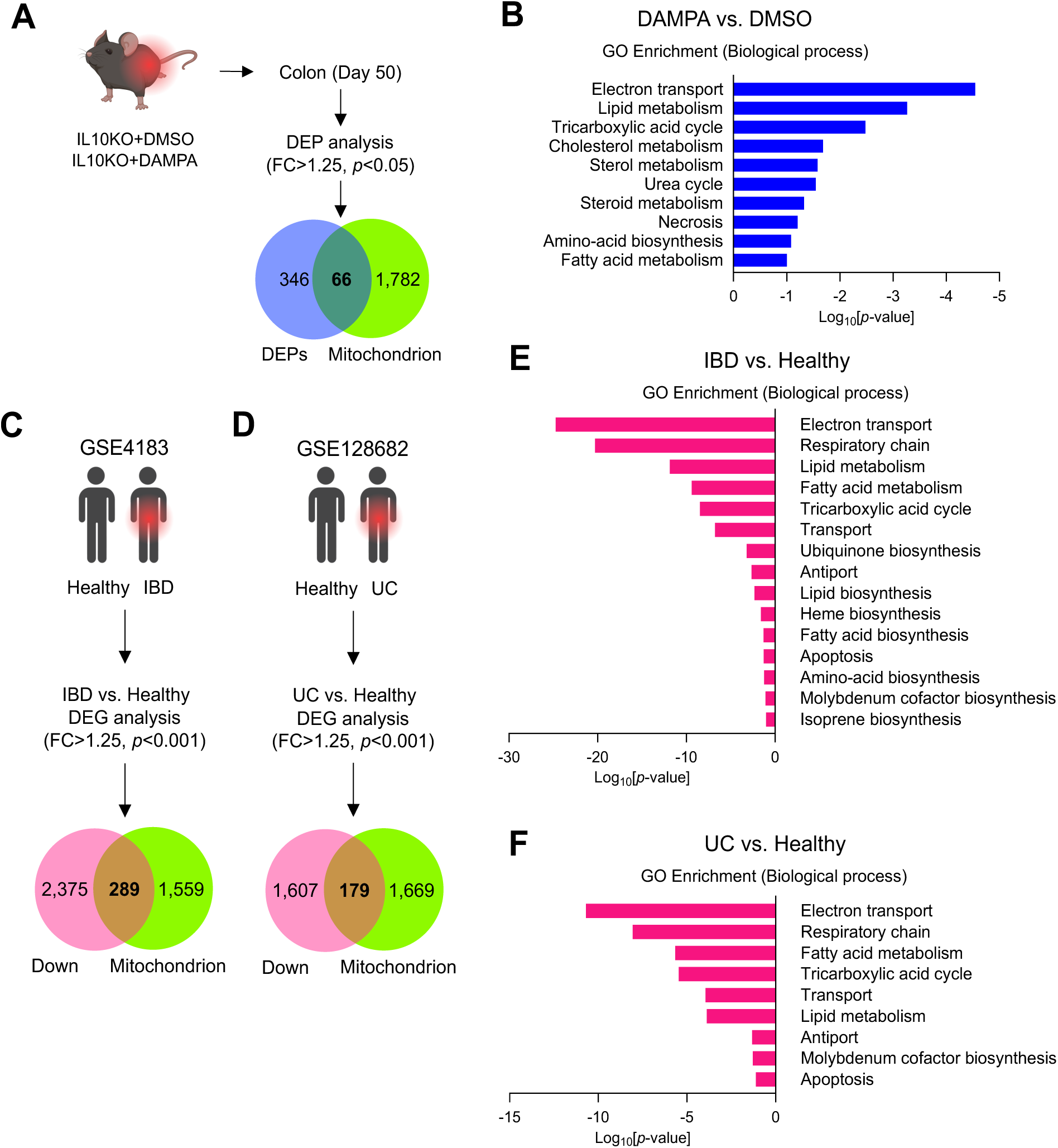
Mitochondrial pathways enriched by DAMPA are suppressed in human IBD. (A and B) Differential protein expression analysis of colonic tissues from IL10KO mice treated with DAMPA for 50 days (related to Figure 6). Differentially expressed proteins (DEPs; cutoff: fold change > 1.25, *p* < 0.05) were identified and intersected with proteins annotated to the Gene Ontology (GO) term “mitochondrion” (GO:0005739). The 66 overlapping proteins were subjected to GO biological process enrichment analysis. (C-F) Differential gene expression and mitochondrial pathway analyses in human IBD datasets. Publicly available transcriptomic datasets comparing intestinal biopsy samples from patients with IBD (GSE4183) or ulcerative colitis (UC; GSE128682) to healthy controls were analyzed (DEG cutoff: fold change > 1.25, *p* < 0.001). (C and E) The 289 mitochondrial genes downregulated in IBD were analyzed for GO biological process enrichment. (D and F) The 179 mitochondrial genes downregulated in UC were analyzed for GO biological process enrichment. Bar graphs show significantly enriched biological processes.

To determine whether similar mitochondrial metabolic pathways are disrupted in human IBD, we analyzed two publicly available transcriptomic datasets (Figures 7C and 7D).^84,85^ GO biological process enrichment analyses of mitochondrial genes significantly downregulated in patients in both datasets also identified the electron transport chain as the most disrupted pathway, followed by lipid metabolism pathways and the TCA cycle (Figures 7E and 7F). This indicates that mitochondrial metabolic dysfunction signatures associated with IBD are restored in DAMPA-treated IL10KO mice, suggesting that DAMPA promotes recovery of mitochondrial metabolism and energetics under inflammatory stress. These findings corroborate the mitoprotective and anti-inflammatory properties of DAMPA observed *in vitro* and *in vivo*.

## DISCUSSION

This study uncovers a previously unrecognized role for gut microbiota-derived MTX metabolite DAMPA in improving intestinal health. We identify gut commensals that produce DAMPA and elucidate its consequence on the intestinal epithelium. Our findings highlight mitochondria as a central signaling hub through which DAMPA modulates epithelial homeostasis during inflammation via STAT3 signaling. Together, these results reframe the role of the gut microbiota in MTX therapy from passive drug inactivation to active regulation of host signaling pathways.

Patients with IBD show highly variable response to MTX therapy.^86^ Gut microbial metabolism of MTX is a critical determinant of therapeutic outcome.^23,87,88^ We established CPDG2 as a key microbial enzyme for DAMPA production in the gut. To test whether intestinal inflammation impairs microbial MTX metabolism to DAMPA, we measured CPDG2 abundance in IBD. Strikingly, we found that CPDG2 abundance is significantly reduced in both the IL10KO IBD mouse model and human IBD cohorts, underscoring the clinical significance of microbiome-mediated DAMPA production. Importantly, inflammation-associated loss of CPDG2 corresponded with a marked reduction in luminal DAMPA production in IL10KO mice. Thus, by defining the microbiome contributors and host effects of MTX metabolism, this work provides a foundation for considering microbiome-informed approaches to improve MTX safety and efficacy in inflammatory diseases.

A previous study has reported a positive correlation between Firmicutes abundance and fecal DAMPA levels.^89^ Our data suggest that inflammation-associated loss of Firmicutes likely contributes to diminished MTX metabolism, supporting a phylum-level contribution to DAMPA production. Consistent with this, we observed reduced Firmicutes abundance in both human IBD cohorts and IL10KO mice, paralleling the decline in CPDG2 and DAMPA production. At the species level, using a combination of *in silico*, *in vitro*, and *in vivo* analyses, we identified *C. asparagiforme* as a potent DAMPA-producing gut bacterium, generating measurable DAMPA within one hour of MTX administration. A prior study reported DAMPA production by *C. symbiosum* at low levels and no detectable DAMPA production by *C. asparagiforme* under comparable MTX concentrations, albeit after 72 h of exposure time.^23^ Our western blot analysis confirmed that both *C. asparagiforme* and *C. symbiosum* express CPDG2. However, in one-hour acute MTX exposure condition, substantial DAMPA production was observed only in *C. asparagiforme*. These discrepancies likely reflect variations in experimental design, including culture conditions and, critically, incubation time. CPDG2 has been reported to deconjugate MTX within minutes, with measurable DAMPA observed as early as 15 minutes after exposure in human subjects.^90^ Our abbreviated timeframe, therefore, provides a physiologically faithful representation of rapid microbial MTX metabolism in the gut. Importantly, growth curve analyses indicate that *C. asparagiforme* retains growth capacity across a broad range of MTX concentrations,^91^ suggesting that MTX does not substantially impair bacterial viability and thus is unlikely to restrict DAMPA production. Whether *C. asparagiforme* further metabolizes or degrades DAMPA *in vitro* after longer incubation times is unknown.

Levels of DAMPA were not completely abolished in IL10KO mice, exhibiting ∼20% of wild-type levels despite no detectable CPDG2 expression. This suggests that additional bacterial enzymes may contribute to DAMPA production. Other hydrolases, such as p-aminobenzoyl-glutamate hydrolase (PGH),^92^ may also convert MTX to DAMPA. Consistently, in our gnotobiotic experiment wherein animals were colonized with a defined community of bacteria with or without *C. asparagiforme*, some DAMPA production was observed in control mice. Inclusion of *C. asparagiforme* led to a modest increase in DAMPA production *in vivo* as compared to our *in vitro* results. Because Firmicutes are known scavengers that rely on products and precursors from other gut commensals for their metabolic capacity,^93^ *C. asparagiforme* may require cooperative metabolic inputs from other commensal gut bacteria to efficiently produce DAMPA *in vivo*. Whether collaborative metabolism of MTX by *C. asparagiforme* and other Firmicutes bacteria enhances DAMPA production in the gut warrants further investigation.

On the host side, we showed that DAMPA improves mitochondrial function and morphology under inflammatory conditions. Mitochondrial dysfunction is increasingly recognized as a central feature of IBD pathogenesis.^94^ Therapeutic strategies aimed at restoring mitochondrial integrity have emerged as potential approaches to limit intestinal inflammation.^95^ Mitochondrial localization of STAT3 (mitoSTAT3) is an important regulator of inflammatory responses.^96^ In this context, our findings suggest that the gut microbial MTX metabolite DAMPA supports mitochondrial resilience under inflammatory conditions through mitoSTAT3 signaling. This work thus highlights a previously unappreciated host–microbe–drug axis linking bacterial metabolism to epithelial mitochondrial health.

We found that DAMPA promotes mitochondrial health by inducing mitophagy in gut epithelial cells, with a concomitant decrease in the mitochondrial fission protein, Dynamin-Related Protein 1 (DRP1). Although an increase in DRP1 is canonically linked to mitophagy, accumulating evidence indicates that mitophagy can proceed independently of DRP1-mediated fragmentation.^83,97,98^ For example, FUNDC1-dependent mitophagy has been shown to occur in the absence of mitochondrial fission, and FUNDC1 signaling can suppress DRP1 interaction and mitochondrial fragmentation.^83,97^ Consistent with these findings, a reduction in DRP1 observed following DAMPA treatment suggests that DAMPA may promote a quality-control mitophagy program that limits excessive mitochondrial fragmentation while preserving mitochondrial network integrity. *In vivo* analyses revealed increased FUNDC1 abundance in colon tissues from DAMPA-treated mice, further supporting a potential role for DAMPA in mitochondrial quality-control pathways.

Our findings suggest that DAMPA induces mitoSTAT3 signaling in gut epithelial cells through activation of the Folate Receptor Alpha (FRα) at the cell surface. Although MTX exhibited lower predicted binding affinity to FRα than DAMPA in our molecular docking analysis, prior studies have reported that MTX can activate nuclear STAT3 signaling, consistent with our observation of increased nuclear p-STAT3. MTX has been shown to elevate pro-inflammatory cytokines such as IL-6 and TNFα in intestinal epithelial cells, which can in turn activate the JAK/STAT3 pathway.^99^ Thus, MTX-induced STAT3 activation may occur indirectly through cytokine-mediated signaling. In contrast, because DAMPA remains luminal and does not enter epithelial cells, its effects on STAT3 are unlikely to arise intracellularly. While future studies using genetic or pharmacologic disruption of FRα will be required to definitively establish FRα-dependent signaling in DAMPA-mediated STAT3 modulation, our findings indicate that MTX and DAMPA may differentially regulate STAT3 through distinct mechanisms.

Structurally, DAMPA differs from MTX in that it lacks the terminal glutamate moiety following CPDG2-mediated cleavage. Considering that the major folate transporters Reduced Folate Carrier (RFC) and Proton-Coupled Folate Transporter (PCFT) require a glutamate tail for efficient substrate recognition and translocation,^100,101^ DAMPA is unlikely to undergo carrier-mediated uptake through these transporters. Consistent with this, we did not detect DAMPA in the intestinal epithelial cell lines, mouse intestinal tissues, or systemic circulation following MTX or DAMPA treatment. A previous study detected DAMPA in serum following high-dose MTX administration (50 mg/kg).^23^ Given that high-dose MTX has been associated with intestinal barrier disruption,^102^ it is possible that the systemic appearance of DAMPA under those conditions reflects passive translocation secondary to epithelial injury rather than carrier-mediated epithelial transport. Notably, accumulating evidence from multiple inflammatory disease models indicates that mitoSTAT3 activation enhances mitophagy, preserves mitochondrial integrity, and attenuates pathogenic cytokine responses.^103,104^ In rheumatoid arthritis and systemic sclerosis models, mitoSTAT3 has been shown to limit inflammatory progression through mitochondrial quality control mechanisms.^103,104^ These findings raise the possibility that, if present systemically, DAMPA-induced mitoSTAT3 signaling could exert an anti-inflammatory effect beyond the intestine. Collectively, these findings expand our understanding of gut microbial metabolite signaling and position microbial MTX metabolism as an active regulator of host mitochondrial and inflammatory homeostasis, with potential implications for microbiome-informed modulation of intestinal inflammation.

## RESOURCE AVAILABILITY

### Lead contact

Further information and requests for information and resources should be directed to and will be fulfilled by the lead contact, Snehal N. Chaudhari.

### Materials availability

This study did not generate new unique reagents.

### Data and code availability

- This paper analyzes existing, publicly available datasets. The accession numbers for human fecal metagenomic sequencing data and human intestinal biopsy transcriptomic data are listed in the Key Resources Table. ShortBRED outputs were processed with custom scripts available at https://github.com/aswalker-lab/CPDG2_metagenomics. ShortBRED markers used are also available in this repository.
- The mass spectrometry proteomics data have been deposited to the ProteomeXchange Consortium via the PRIDE^105^ partner repository with the dataset identifier PXD075428 for human HT29 proteomics data and PXD075463 for the mouse colon tissue proteomics data. Proteomics data are publicly available as of the date of publication.
- Microbiome 16s rRNA sequencing data have been deposited in the NCBI Sequence Read Archive (SRA) database under accession number SRA: PRJNA1432080. Data are publicly available as of the date of publication.
- All ImageJ macros used for MiNA analysis and colocalization quantification have been deposited at GitHub and are publicly available at https://github.com/krw-uw/modified-MiNA-macro-for-mitochondrial-analysis.
- All other data reported in this paper will be shared by the lead contact upon request.

## ACKNOWLEDGEMENTS

This study was supported by the National Institutes of Health R00 DK128503 (S.N.C), the Ruth Dickie Endowment from the Madison Chapter of the Graduate Women in Science (J.C.), National Institutes of Health T32 GM135066 (J.W.), T32 GM152341 (W.S.K.), R35 GM146987 (A.S.W.), the Wisconsin Alumni Research Foundation (WARF), and the department of Biochemistry at University of Wisconsin-Madison. We thank Dr. Kurt Weiss for assistance with fluorescence imaging and mitochondrial network analysis. We thank the UWCCC Experimental Animal Pathology Laboratory for use of its facilities and services. We thank Drs. Eric Yen, Ting Fu, and Margaret Alexander for insightful guidance that strengthened this work. We also acknowledge the Nutrition and Metabolism PhD Program and the Department of Biochemistry at the University of Wisconsin–Madison for their institutional support. Finally, we thank members of the Chaudhari lab for technical assistance and helpful discussions.

## AUTHOR CONTRIBUTIONS

Conceptualization, J.C., and S.N.C.; Methodology, J.C, J.W., and S.N.C.; investigation, J.C., R.D., W.S.K., A.S., J.A.H., J.V., A.S.W. and S.N.C.; writing-original draft, J.C. and S.N.C.; writing-review & editing, J.C., R.D., W.S.K., J.W., A.S., J.A.H., J.V., A.S.W., T.R., and S.N.C.; funding acquisition, J.C., and S.N.C.; resources, A.S., J.V., A.S.W., and T.R.; supervision, S.N.C.

## SUPPLEMENTAL INFORMATION

Document S1. Figures S1–S11

Table S1. Phylum-level changes in IBD patients. Related to Figure 2.

Table S2. The primer list for qPCR. Related to STAR Methods.

## STAR★METHODS

KEY RESOURCES TABLE.

**KEY RESOURCES TABLE**
| REAGENT or RESOURCE | SOURCE | IDENTIFIER |
| --- | --- | --- |
| **Antibodies** |  |  |
| Rabbit CPG2 Antibody | Aviva Systems Biology | Cat #OACA04573, RRID: AB_3739805 |
| Rabbit LC3B Antibody | Cell Signaling Technology | Cat #2775, RRID: AB_915950 |
| Rabbit PINK1 Polyclonal Antibody | Invitrogen | Cat #PA1-16604, RRID: AB_2164267 |
| Rabbit FUND1 Antibody (N-term) | Abcepta | Cat #AP17377a RRID: AB_11134968 |
| Rabbit Phospho-STAT3 (phospho Tyr705) Antibody | GeneTex | Cat #GTX118000, RRID: AB_10618482 |
| Rabbit Phospho-STAT3 (phospho Ser727) Antibody | Cell Signaling Technology | Cat #9134, RRID: AB_331589 |
| Rabbit β-Actin Antibody | Cell Signaling Technology | Cat #8457, RRID: AB_10950489 |
| Mouse anti-rabbit IgG-HRP | Santa Cruz Biotechnology | Cat #sc-2357, RRID: AB_628497 |
| Goat Anti-Rabbit IgG (H+L) Antibody, Alexa Fluor 488 Conjugated | Invitrogen | Cat #A-11008, RRID: AB_143165 |
| **Bacterial strains** |  |  |
| *Akkermansia muciniphila* | Clardy lab at Harvard Medical School | N/A |
| *Bifidobacterium longum subspecies infantis* | ATCC | ATCC 15697 |
| *Bacteroides thetaiotaomicron* | Devlin lab at Harvard Medical School | N/A |
| *Clostridium scindens* | Devlin lab at Harvard Medical School | N/A |
| *Clostridium asparagiforme* | Rey lab at the University of Wisconsin-Madison | N/A |
| *Clostridium symbiosum* | ATCC | ATCC 14940 |
| *Escherichia coli* Nissle 1917 | Lesser lab at Tufts University | N/A |
| *Fusobacterium nucleatum* subsp. *nucleatum* | ATCC | ATCC 25586 |
| **Chemicals, peptides, and recombinant proteins** |  |  |
| Methotrexate (MTX) | Schircks Laboratories | Cat #16.411 |
| Methotrexate disodium salt | Thermo Scientific Chemicals | Cat #AAJ66364MD |
| 4-Amino-10-methylpteroic acid (DAMPA) | Schircks Laboratories | Cat #16.403 |
| Tumor Necrosis Factor-alpha (TNFα) human | Sigma Aldrich | Cat #H8916 |
| Folic acid-d2 | Santa Cruz Biotechnology | Cat #sc-489921 |
| Rotenone | Sigma Aldrich | Cat #R8875 |
| Oligomycin A | Sigma Aldrich | CAS #79-13-5 |
| FCCP | Sigma Aldrich | Cat #C2920-10MG |
| Antimycin A | Fisher Scientific | Cat #NC0862023 |
| Stattic | MedChemExpress | Cat #HY-13818 |
| **Critical commercial assays** |  |  |
| No-Stain Protein Labeling Reagent | Invitrogen | Cat #A44449 |
| ZymoBIOMICS 96 MagBead DNA Kit | Zymo Research | Cat #D4302 |
| ZymoBIOMICS DNA Miniprep Kit | Zymo Research | Cat #D4300 |
| Quick-16S NGS Library Prep Kit | Zymo Research | Cat #D6400 |
| Mouse Lipocalin-2/NGAL DuoSet ELISA | R&D Systems | Cat #DY1857 |
| MitoTracker™ Red CMXRos | Invitrogen | Cat #M7512 |
| LysoTracker™ Green | Cell Signaling Technology | Cat #8783S |
| ER-Tracker™ Red | Invitrogen | Cat #E34250 |
| CellTiter-Glo® Luminescent Cell Viability Assay | Promega | Cat #G7571 |
| MitoSOX™ Red Mitochondrial Superoxide Indicator | Invitrogen | Cat #M36008 |
| Maxima First Strand cDNA Synthesis Kit | Thermo Scientific | Cat #K1671 |
| LightCycler 480 SYBR Green | Roche | Cat #04707516001 |
| Seahorse Xfe24 Fluxpak Mini | Agilent Technologies | Cat #102342-100 |
| CYTO-ID® Autophagy Detection Kit 2.0 | Enzo Life Sciences | Cat #KIT175 |
| Lipofectamine™ 3000 Transfection Reagent | Invitrogen | Cat #L3000150 |
| STAT3 Reporter Kit | BPS Bioscience | Cat #79730 |
| Lipofectamine 3000 Transfection Reagent | Invitrogen | Cat #L3000150 |
| Dual-Luciferase® Reporter Assay System | Promega | Cat #E1910 |
| LEGENDplex™ Mouse Anti-Virus Response Panel (13-plex) with V-bottom Plate | BioLegend | Cat #740622 |
| **Deposited data** |  |  |
| Metagenome sequencing data | Franzosa et al.<sup>29</sup><br>Zuo et al.<sup>30</sup> | SRA: PR000677<br>SRA: PRJNA759642 |
| RNA sequencing raw data | Fenton et al.<sup>84</sup> | GSE128682, GSE4183 |
| Proteomics | This study | PXD075428, PXD075463 |
| Microbiome sequencing data | This study | SRA: PRJNA1432080 |
| **Experimental models: Cell lines** |  |  |
| *Homo sapiens*, HT29 | ATCC | ATCC HTB-38 |
| *Homo sapiens*, Caco-2 | ATCC | ATCC HTB-37 |
| **Experimental models: Organisms/strains** |  |  |
| Mouse: C57BL/6J | Jackson Laboratory | RRID:IMSR_JAX:000664 |
| Mouse: IL10KO | Jackson Laboratory | RRID:IMSR_JAX:002251 |
| Mouse: Germ-free | National Gnotobiotic Rodent Resource Center at the University of North Carolina at Chapel Hill | N/A |
| **Oligonucleotides** |  |  |
| Primers for qPCR | Table S2 | N/A |
| **Software** |  |  |
| ImageJ 1.54p | NIH | https://imagej.net/software/fiji/ |
| GraphPad Prism v10.0 | GraphPad | https://graphpad.com/ |
| QuPath v0.6.0 | Bankhead, P. et al.<sup>106</sup> | https://qupath.github.io/ |
| Metascape | Zhou et al.<sup>44</sup> | https://metascape.org/ |
| Agilent Seahorse Wave Controller Software v2.6.4.24 | Agilent | https://www.agilent.com/en/support/seahorse-xf-software-releases |
| FreeStyle | Thermo Fisher | N/A |
| Neurosnap | Neurosnap Inc.<sup>107</sup> | https://neurosnap.ai/ |
| Protein BLAST | NIH | https://blast.ncbi.nlm.nih.gov/Blast.cgi |
| Attune NxT software | Thermo Fisher | N/A |
| NIS Elements software | Nikon | N/A |
| PyMOL 3.1.6.1 | Schrödinger, LLC.<sup>108</sup> | https://www.pymol.org/ |
| ChemDraw v22.0.0 | PerkinElmer | N/A |
| Biorender | Biorender | https://biorender.com/ |
| **Other** |  |  |
| 12 mm Transwell® with 0.4 µm Pore Polycarbonate Membrane Insert | Corning | Cat #3401 |
| Corning® 24 mm Transwell® with 8.0 µm Pore Polycarbonate Membrane Insert | Costar | Cat #3428 |
| Cell Lysis Buffer | Cell Signaling Technology | Cat #CSIG-9803S |
| cOmplete™ Protease Inhibitor Cocktail | Roche | Cat #11836170001 |
| Bradford Dye Reagent | Thermo Scientific Chemicals | Cat #J61522.AP |
| Bovine Serum Albumin | Tocris | Cat #5217 |
| SuperSignal™ West Pico Plus Chemiluminescent Substrate | Thermo Scientific | Cat #PI34577 |
| StartingBlock™ Blocking Buffer | Thermo Scientific | Cat #37538 |
| Bolt™ Bis-Tris Plus Mini Protein Gels, 4-12%, 10-well | Invitrogen | Cat #NW04122BOX |
| Bolt™ Bis-Tris Plus Mini Protein Gels, 4-12%, 15-well | Invitrogen | Cat #NW04125BOX |
| PVDF/Filter Paper Sandwiches | Invitrogen | Cat #LC2005 |
| NuPAGE™ MES SDS Running Buffer | Invitrogen | Cat #NP0002 |
| Bolt™ Transfer Buffer | Invitrogen | Cat #BT00061 |
| Tris Buffered Saline with Tween 20 | Santa Cruz | Cat #sc-281695 |
| Strata-X 33 µm Polymeric Reversed Phase | Phenomenex | Cat #8E-S100-TGB |
| Luna Omega 1.6µm Polar C18 Column | Phenomenex | Cat #00D-4748-AN |
| Seahorse XFe24 FluxPak mini | Agilent | Cat #102342-100 |
| Seahorse XF DMEM medium, pH 7.4 | Agilent | Cat #103575-100 |
| Seahorse XF 1M Glucose solution | Agilent | Cat # 103577-100 |
| Seahorse XF 100 mM Pyruvate solution | Agilent | Cat # 103578-100 |
| Seahorse XF 200 mM Glutamine solution | Agilent | Cat #103579-100 |
| TRIzol™ Reagent | Invitrogen | Cat #15596026 |
| Hoechst 33342 | Thermo Scientific | Cat #62249 |
| Donkey serum | GeneTex | Cat #GTX27475 |
| TMB Substrate | R&D Systems | Cat #DY999B |
| Stop Solution | R&D Systems | Cat #DY994 |

## KEY RESOURCES TABLE

## EXPERIMENTAL MODEL AND STUDY PARTICIPANT DETAILS

### Mice

C57BL/6J male mice (9-11 weeks old) and IL10 knockout (IL10KO) male mice (9-11 weeks old; B6.129P2-Il10tm1Cgn/J) were purchased from Jackson Laboratory and bred in the University of Wisconsin-Madison Biomedical Research Model Services (BRMS) facility. Animals are transferred to the Chaudhari lab animal facility for experiments. Germ-free male mice (6-8 weeks old) were obtained from the National Gnotobiotic Rodent Resource Center (NGRRC) at the University of North Carolina at Chapel Hill and bred in the University of Wisconsin-Madison BRMS Gnotobiotic Shared Resource (GSR) facility. Germ-free animals were maintained in a controlled environment in plastic flexible film gnotobiotic isolators.

For short-term MTX exposure experiments, WT and IL10KO mice received a single oral gavage of MTX-Na (5 mg/kg) dissolved in sterile water and euthanized 1 h after MTX administration. For the chronic DAMPA treatment, IL10KO mice were orally gavaged daily with DAMPA (4 mg/kg) or vehicle for 50 consecutive days. DAMPA was dissolved in 10% DMSO in sterile water, and control animals received 10% DMSO in sterile water.

For germ-free colonization studies, animals were colonized with a defined bacterial consortium either containing or lacking *C. asparagiforme* by oral gavage two weeks prior to MTX treatment. On the day of the experiment, mice received MTX-Na (5 mg/kg) by oral gavage, and cecum contents were harvested 1 h after MTX administration for 16S rRNA sequencing and DAMPA quantification via LC-MS.

All animals were housed in climate-controlled animal facilities under a 12 h light/12 h dark cycle with ad libitum access to food and water. Mice were fasted for 4 h prior to sacrifice. Tissues, intestinal contents, and fecal samples were immediately snap-frozen on dry ice and stored at –80°C until further analysis. For serum, blood was allowed to clot at room temperature for 30 min and centrifuged at 1,000 x g for 20 min. The supernatant was transferred to microcentrifuge tubes and stored at −80°C until further analysis. All procedures were approved by the Institutional Animal Care and Use Committee (IACUC) at the University of Wisconsin-Madison.

### Bacteria

Information for the eight bacterial strains used in this study is provided in the Key Resources Table. Bacteria were cultured anaerobically in Cullen-Haiser Gut (CHG) medium,^109^ consisting of brain heart infusion (BHI) medium (Dot Scientific) supplemented with 1% BBL vitamin K_1_-hemin solution (BD), 1% trace minerals solution (ATCC), 1% trace vitamins solution (ATCC), 5% fetal bovine serum (FBS, GenClone), 1 g/L cellobiose, 1 g/L maltose, 1 g/L fructose, and 0.5 g/L cysteine. Cultures were maintained in an anaerobic chamber (Coy Laboratory Products) with a gas mixture of 5% H_2_, 20% CO_2_, and balance N_2_ at 37°C.

For western blot analysis, 100 µL of overnight cultures were inoculated into 3 mL of reduced CHG medium and incubated anaerobically at 37°C overnight. For DAMPA production measurements, 2 mL of overnight cultures were diluted two-fold in fresh CHG medium, supplemented with 200 µM MTX-Na and 2 mM ZnCl_2_, and incubated for 1 h under anaerobic conditions. Cultures were then tightly sealed in 15 mL conical tubes before removal from the anaerobic chamber to minimize oxygen exposure and immediately centrifuged at 4,000 x g for 5 min. Cell pellets were collected and processed for further analyses.

### Intestinal cell cultures

HT29 cells and Caco-2 cells were obtained from ATCC. Cells were cultured in Dulbecco’s Modified Eagle Medium (DMEM, Gibco), supplemented with 10% FBS (GenClone) and 1% penicillin and streptomycin (GenClone) in a humidified atmosphere at 37°C with 5% CO_2_. Cells were split every three to four days using 0.25% Trypsin-EDTA (Santa Cruz Biotechnology) and used in experiments two to three days after plating.

For DAMPA and MTX treatment, HT29 cells were pre-treated with TNFα (10 ng/mL) for 24 h, followed by co-treatment with 25 µM DAMPA or 25 µM MTX for an additional 24 h in the continued presence of TNFα. DAMPA and MTX were dissolved in DMSO, and the final DMSO concentration was 0.5%. Vehicle controls received an equivalent concentration of DMSO. All treatments were performed in complete growth medium.

## METHOD DETAILS

### Preparation of bacterial consortium

Six bacterial species (*B. longum, B. thetaiotaomicron, C. asparagiforme, F. nucleatum, E. coli Nissle 1917*, and *A. muciniphila*) were grown anaerobically in CHG medium. Each mouse was gavaged with 200 µL of the bacterial community, consisting of ∼2.4 x 10^8^ cells per species per dose. Bacterial mixtures were prepared fresh and transferred in tightly sealed tubes to minimize oxygen exposure prior to gavage. Cell density was estimated using McFarland standards as an approximate measure of bacterial concentration. McFarland unit (MFU) 10 corresponds to approximately 3 x 10^9^ cells per mL.^110^

### Transwell-based transport assays

HT29 and Caco-2 cells were cultured on permeable polycarbonate membrane inserts in Transwell plates and maintained at 37°C with 5% CO2 in complete growth medium, which was replaced every two days. Experiments were performed after formation of a confluent monolayer, approximately 3 weeks post-seeding. HT29 cells were seeded onto 8 µm pore-size inserts in 6-well plates at a density of 200,000 cells per insert. Cells were treated with 25 µM MTX or 25 µM DAMPA for 1 h. Vehicle controls received 0.5% DMSO. Following treatment, cells were washed twice with PBS, scraped, and centrifuged at 250 x g for 5 min at 4°C. Pellets were stored at −80°C for subsequent LC-MS analysis. Caco-2 cells were seeded onto 0.4 µm pore-size inserts in 24-well plates at a density of 8.5 x 10^4^ cells per 0.33 cm^2^ insert. After monolayer formation, cells were treated with 25 µM MTX or DAMPA for 24 h. Apical and basolateral medium (100 µL) were collected and stored at −80°C for LC-MS analysis.

### Western blot

Mouse colon tissues, fecal samples, and bacterial cell pellets were lysed in 200-300 µL of ice-cold 1x Cell Lysis Buffer (Cell Signaling Technology) supplemented with protease inhibitor cocktail (Roche) and homogenized using a Bead Ruptor Elite (Omni International). Samples were centrifuged at 18,200 x g for 10 min at 4°C, and supernatants were collected. Protein concentrations were determined using a Bradford assay (Thermo Scientific) with bovine serum albumin (BSA) as a standard. Proteins were mixed with 4x loading buffer supplemented with β-mercaptoethanol, boiled at 100°C for 10 min, and separated on precast Bis-Tris SDS-PAGE gels (Invitrogen). Proteins were transferred onto PVDF membranes using wet transfer at 30 V for 1 h. Membranes were blocked in StartingBlock™ Blocking Buffer (Thermo Scientific) for 1 h at room temperature and incubated with primary antibodies diluted in 3% BSA in TBS-T overnight at 4°C. After washing with TBS-T, membranes were incubated with HRP-conjugated secondary antibodies diluted in 3% BSA in TBS-T for 1 h at room temperature. Blots were developed using SuperSignal™ West Pico PLUS Chemiluminescent Substrate (Thermo Scientific). Blot images were obtained from iBright imaging system (Invitrogen). Band intensities were quantified using the iBright imaging system (Invitrogen) with automated band detection and normalized to total protein or β-actin. Primary antibodies used for western blot are: CPDG2 (1:1,000; Aviva Systems Biology), PINK1 (1:1,000; Invitrogen), β –actin (1:1,000; Cell Signaling), and LC3B (1:500; Cell Signaling). Mouse anti-rabbit IgG-HRP (1:10,000; Santa Cruz) was used as a secondary antibody.

### Metabolite extraction and LC-MS analysis

Extraction and quantification of MTX and DAMPA were performed using a previously established method optimized for folate metabolites.^28^

For HT29 and Caco-2 cell pellets, samples were resuspended in 200 µL of freshly prepared folate extraction buffer (FEB; 20 mM ascorbic acid, 20 mM ammonium acetate, 20 mM β-mercaptoethanol (BME), pH 8.1 adjusted with NaOH) and transferred into 2 mL tubes with glass beads. Cecum contents (50-100 mg) were processed similarly in 400 µL FEB with glass beads. Samples were homogenized for 1 min at 4 m/s using a Bead Ruptor Elite (Omni International). Apical and basolateral medium (100 µL) were mixed with 300 µL FEB without homogenization.

For bacterial cell pellets, 200 µL of extraction solvent (50:50 methanol:water, 20 mM ascorbic acid, 20 mM BME) was added, vortexed for 30 sec to lyse cells, and centrifuged at 18,200 x g for 5 min. Supernatants were dried in a SpeedVac (Thermo Scientific), and reconstituted in 200 µL FEB.

Following sample-specific extraction, rat serum (5 µL per 200 µL FEB) was added to each sample, followed by incubation at 37°C for 45 min for deconjugation. Samples were subsequently heated at 100°C for 5 min and immediately cooled on ice. After centrifugation at 18,200 x g for 5 min at 4°C, the supernatants were subjected to solid-phase extraction (SPE) using Strata-X columns (Phenomenex, #8E-S100-TGB). Columns were conditioned with 200 µL of methanol and equilibrated with 200 µL of resuspension buffer (20 mM ammonium acetate, 20 mM BME) prior to sample loading. Supernatants were applied to the SPE columns and washed with 200 µL of resuspension buffer. Analytes were eluted using 200 µL of elution buffer (50:50 methanol:water, 20 mM ammonium acetate, 20 mM BME). Eluates were dried in a SpeedVac (Thermo Scientific), reconstituted in 40 µL of resuspension buffer, and used for LC-MS analysis.

LC-MS analysis was performed on a Vanquish ultra-high-performance liquid chromatography system (Thermo Scientific) coupled to an Orbitrap Exploris 120 mass spectrometer (Thermo Scientific). Chromatographic separation was achieved on a reversed-phase polar C18 column (Phenomenex, #00D-4748-AN) maintained at 40°C using a 12-minute gradient with solvent A (95% H_2_O, 5% methanol, 5 mM dimethyl-hexylamine (DMHA), pH 8.1 adjusted with formic acid) and solvent B (100% methanol, 5 mM DMHA) at a flow rate of 0.4 mL/min. Eluted metabolites were analyzed by heated electrospray ionization in negative mode. Quantification was performed by integrating peak areas using Freestyle software. Signals were normalized to the internal standard (folic acid-d_2_) and to sample weight for tissue-derived samples. Concentrations were calculated using linear regression based on standard curves.

### Human metagenomic data analysis

Human metagenomics data was downloaded from NCBI using SRA tools. A representative fastq file for each dataset file was checked for quality and adapters using FastQC (https://www.bioinformatics.babraham.ac.uk/projects/fastqc/). The representative file from Fransoza et al showed significant presence of the Universal adapter. All fastq files from this dataset were therefore trimmed using trimmomatic (https://github.com/usadellab/Trimmomatic).^111^ We then used shortBRED (https://huttenhower.sph.harvard.edu/shortbred/)^112^ to quantify the abundance of reads that mapped onto CPDG2. shortBRED markers were constructed from CPDG2 sequences of *C.asparagiforme*, *C. symbiosum*, and *C. innocuum* identified through BLAST search on NCBI along with additional Uniprot sequences annotated as Carboxypeptidase G2. Markers and the fasta file used to generate the markers are available in the Data and code availability section. ShortBRED was then used to quantify marker abundance across metagenomic sequencing data. Total abundance across all markers was calculated and fastq files were matched to disease state using the study metadata.

### Sequence similarity analysis of CPDG2

To screen CPDG2 protein sequence similarity across 43 gut bacterial species, CPDG2 (UniProt accession: P06621) was used as a query for protein-protein BLAST (BLASTP) searches using the NCBI BLAST against the ClusteredNR database. Word size was set to 6, and default scoring parameters were applied. For each bacterial species, the best hit based on the lowest E-value (expectation value) was selected, and the corresponding maximum alignment score was used for comparative analysis and visualization in a heatmap.

### 16S rRNA sequencing

16S rRNA gene sequencing was performed by Zymo Research using the ZymoBIOMICS^®^ Targeted Sequencing Service. Genomic DNA was extracted using the ZymoBIOMICS^®^-96 MagBead DNA Kit or ZymoBIOMICS® DNA Miniprep Kit according to the manufacturer’s protocols. The V3–V4 region of the bacterial 16S rRNA gene was amplified and libraries were prepared using the Quick-16S™ NGS Library Prep Kit (Zymo Research). Libraries were sequenced on an Illumina NextSeq platform (600-cycle kit). Amplicon sequence variants (ASVs) were inferred using the DADA2 pipeline (Callahan et al.^113^), and chimeric sequences were removed during processing. Taxonomic assignment and diversity analyses were performed using QIIME v1.9.1 (Caporaso et al.^114^) with the Zymo Research reference database.

### Untargeted proteomics

#### Sample preparation for mass spectrometry

Samples for protein analysis were prepared essentially as previously.^115,116^ HT29 cells were washed three times with ice-cold PBS. 100 µL of lysis buffer (8 M urea, 200 mM EPPS pH 8.5, cOmplete™ Protease Inhibitor Cocktail (Roche), 0.1% SDS) was added and shaken for 10 min at 4°C to allow cell lysis. 100 µL of nuclease buffer (200 mM EPPS pH 8.5, 0.5 µL/mL Pierce™ Universal Nuclease (Thermo Fisher)) was then added. For mouse colon tissue samples, tissues were transferred to 2 mL tubes with glass beads containing lysis buffer (8 M urea, 200 mM EPPS pH 8.5, cOmplete™ Protease Inhibitor Cocktail (Roche), 0.1% SDS). Samples were homogenized for 1 min at 4 m/s using a Bead Ruptor Elite (Omni International). To ensure genomic DNA shearing, lysates were passed through a needle ten times. Lysates were transferred to fresh tubes and stored at −80°C until further processing.

Following lysis, 25 μg of protein from each sample was reduced with tris(2-carboxyethyl)phosphine (TCEP) (Thermo Scientific, #77720), alkylated with iodoacetamide (MP Biochemicals, #100351), and then further reduced with dithiothreitol (DTT) (Alfa Aesar, #A15797). A buffer exchange was carried out using a modified SP3 protocol.^117^ Briefly, ∼250 µg of Cytiva SpeedBead Magnetic Carboxylate Modified Particles (65152105050250 and 4515210505250), mixed at a 1:1 ratio, were added to each sample. 100% ethanol was added to each sample to achieve a final ethanol concentration of at least 50%. Samples were incubated with gentle shaking for 15 min. Samples were washed three times with 80% ethanol. Protein was eluted from SP3 beads using 200 mM EPPS pH 8.5 containing Lys-C (Wako, #129-02541). Samples were digested overnight at room temperature with vigorous shaking. The next morning trypsin (1:50 enzyme to protein ratio) (Thermo Scientific, #90305) was added to each sample and further incubated for 6 h at 37° C. Acetonitrile was added to each sample to achieve a final concentration of ∼33%. Each sample was labelled, in the presence of SP3 beads, with ∼62.5 µg of TMTPro reagents (Thermo Fisher Scientific). Following confirmation of satisfactory labelling (>97%), excess TMT was quenched by addition of hydroxylamine to a final concentration of 0.3%. The full volume from each sample was pooled and acetonitrile was removed by vacuum centrifugation for 1 h. The pooled sample was acidified, and peptides were de-salted using a Sep-Pak 50mg tC18 cartridge (Waters). Peptides were eluted in 70% acetonitrile, 1% formic acid and dried by vacuum centrifugation.

#### Basic pH reversed-phase separation (BPRP)

TMT labeled peptides were solubilized in 5% acetonitrile/10 mM ammonium bicarbonate, pH 8.0 and ∼300 µg of TMT labeled peptides were separated by an Agilent 300 Extend C18 column (3.5 µm particles, 4.6 mm ID and 250 mm in length). An Agilent 1260 binary pump coupled with a photodiode array (PDA) detector (Thermo Scientific) was used to separate the peptides. A 45-minute linear gradient from 10% to 40% acetonitrile in 10 mM ammonium bicarbonate pH 8.0 (flow rate of 0.6 mL/min) separated the peptide mixtures into a total of 96 fractions (36 seconds). A total of 96 fractions were consolidated into 24 samples in a checkerboard fashion and vacuum dried to completion. Each sample was desalted via Stage Tips and re-dissolved in 5% formic acid/ 5% acetonitrile for LC-MS3 analysis.

#### Liquid chromatography separation and tandem mass spectrometry (LC-MS3)

Proteome data were collected on an Orbitrap Eclipse mass spectrometer (Thermo Fisher Scientific) coupled to a Proxeon EASY-nLC 1000 LC pump (Thermo Fisher Scientific). A FAIMS device was enabled during data collection with CV values set to –40, –60, and –80. Fractionated peptides were separated using a 120 min gradient at 500 nL/min on a 35 cm column (i.d. 100 μm, Accucore, 2.6 μm, 150 Å) packed in-house. MS1 data were collected in the Orbitrap (60,000 resolution; maximum injection time 50 ms; AGC 4 × 105). Charge states between 2 and 5 were required for MS2 analysis in the ion trap, and a 120 second dynamic exclusion window was used. Top 10 MS2 scans were performed in the ion trap with CID fragmentation (isolation window 0.5 Da; Turbo; NCE 35%; maximum injection time 35 ms; AGC 1 × 104). Real-time search was used to trigger MS3 scans for quantification.^118^ MS3 scans were collected in the Orbitrap using a resolution of 50,000, NCE of 55%, maximum injection time of 250 ms, and AGC of 1.25 × 105. The close out was set at two peptides per protein per fraction.

### Data analysis

Raw files were converted to mzXML, and monoisotopic peaks were re-assigned using Monocle.^119^ Searches were performed using the Comet search algorithm against a human database (May 2021) or mouse database (May 2025) downloaded from Uniprot. A 50 ppm precursor ion tolerance, 1.0005 fragment ion tolerance, and 0.4 fragment bin offset was used for MS2 scans collected in the ion trap. TMTpro on lysine residues and peptide N-termini (+304.2071 Da) and carbamidomethylation of cysteine residues (+57.0215 Da) were set as static modifications, while oxidation of methionine residues (+15.9949 Da) was set as a variable modification.

Each run was filtered separately to 1% False Discovery Rate (FDR) on the peptide-spectrum match (PSM) level. Then proteins were filtered to the target 1% FDR level across the entire combined data set. For reporter ion quantification, a 0.003 Da window around the theoretical m/z of each reporter ion was scanned, and the most intense m/z was used. Reporter ion intensities were adjusted to correct for isotopic impurities of the different TMTpro reagents according to manufacturer specifications. Peptides were filtered to include only those with a summed signal-to-noise (SN) ≥ 180 across all TMT channels. For each protein, the filtered peptide TMTpro SN values were summed to generate protein or phosphorylation site quantification values. The signal-to-noise (S/N) measurements of peptides assigned to each protein were summed (for a given protein). These values were normalized so that the sum of the signal for all proteins in each channel was equivalent thereby accounting for equal protein loading, and *p* values were adjusted for multiple comparisons using the Benjamini-Hochberg (BH) correction.

### Functional enrichment analysis

Gene Ontology (GO) and pathway enrichment analyses were performed using Metascape and DAVID.^44,120,121^ For HT29 proteomics, differentially expressed proteins (DEPs; cutoff: fold-change > 1.25 and *p* value < 0.05) from each comparison (DAMPA vs. DMSO and DAMPA vs. MTX) were used for enrichment analysis. Databases are GO biological process (BP) and GO cellular component (CC), and KEGG immune-related pathways were analyzed using default parameters. For colon tissue proteomics, DEPs (fold change > 1.25 and *p* < 0.05) were intersected with proteins annotated to the GO term “mitochondrion” (GO:0005739), and the overlapping proteins were subjected to GO biological process enrichment analysis. For publicly available transcriptomic datasets (GSE4183 and GSE128682), downregulated genes in human IBD patients (fold change > 1.25 and *p* < 0.001) were intersected with the genes annotated to the GO term “mitochondrion” (GO:0005739), and the overlapping proteins were analyzed for GO biological process enrichment.

### Transmission electron microscopy

HT29 cells were fixed in a modified Karnovsky’s fixative (2.5% glutaraldehyde/2.0% formaldehyde in 0.1 M sodium phosphate buffer (PB), pH 7.2), post fixed in 1.0% osmium tetroxide in 0.1 M PB, dehydrated in a grade series of ethanol, transitioned in acetone, and embedded in EmBed 812 (Electron Microscopy Sciences). Polymerized samples were sectioned (80 nm) on an ultramicrotome and sections were placed onto formvar coated 2×1 Cu slot grids and post-stained with uranyl acetate and lead citrate. The sections were viewed at 80 kV on a FEI CM120 TEM (Thermo Fisher Scientific) and imaged on an AMT BioSprint12 digital camera (AMT Imaging).

HT29 cells grown on coverslips in 24-well plates were processed at the UW-Madison Electron Microscopy Facility. Samples were fixed, embedded, sectioned, and stained according to standard facility protocols. Sections were imaged using an FEI CM120 transmission electron microscope. Mitochondria were classified as normal (lamellar cristae) or abnormal (honeycomb, onion-like, or loss of cristae) based on cristae morphology.^122,123^ The proportion of mitochondrial morphologies was quantified from TEM images. Images acquired at >15,000x magnification were used for cristae morphology assessment, and images at 5600x magnification were used for quantification of autophagic structures. Images were randomly acquired by an independent investigator to minimize region-specific bias.

### Mitochondrial fluorescence staining

Cells were incubated with 100 nM MitoTracker Red CMXRos (Invitrogen) in pre-warmed growth medium at 37°C for 30 min, washed with PBS, and fixed with 4% paraformaldehyde in PBS for 10 min at room temperature. Nuclei were counterstained with DAPI (1:1,000 in PBS) or Hoechst (1:2,000 in PBS) 33342 for 10 min. Fluorescence intensity was measured by a Varioskan™ LUX Multimode Microplate Reader (Thermo Scientific) (MitoTracker: Ex 579 nm/Em 599 nm; DAPI: Ex 358 nm/Em 461 nm), and images were obtained using appropriate filter sets on a Nikon A1R+ confocal microscope (Nikon Instruments) at the Biochemistry Optical Core (BOC) Facility at UW–Madison.

### Lysosome staining and mitochondria–lysosome tethering analysis

Lysosomes were stained using LysoTracker™ Green DND-26 (Cell Signaling Technology). Cells were incubated with 50 nM LysoTracker Green in complete growth medium at 37°C for 30 min and washed once with PBS. Nuclei were counterstained with Hoechst 33342 (1:2,000 in PBS) for 10 min at room temperature. For fluorescence quantification, LysoTracker intensity was measured using a fluorescence microplate reader (LysoTracker: Ex 504 nm/Em 511 nm; Hoechst 33342: Ex 352 nm/Em 455 nm) and normalized to Hoechst fluorescence intensity to account for differences in cell number. Cells were imaged live immediately after staining. Images were obtained using appropriate filter sets on a Nikon A1R+ confocal microscope (Nikon Instruments) at the Biochemistry Optical Core (BOC) Facility at UW–Madison.

For mitochondria–lysosome tethering analysis, cells were co-stained with 100 nM MitoTracker Red CMXRos (Invitrogen) and 50 nM LysoTracker Green in pre-warmed growth medium at 37°C for 30 min, followed by Hoechst 33342 staining (1:2,000 in PBS) for 10 min at room temperature. Live-cell imaging was performed immediately after staining. Mitochondria–lysosome tethering events were quantified by manually counting contact sites and normalizing to the total number of lysosomes per cell. Normalization was performed to lysosome number, as total lysosome abundance was not altered across treatment groups. Image selection was performed by an independent investigator, and quantification was conducted in a blinded manner.

### Endoplasmic reticulum (ER) staining

Cells were incubated with 1 µM ER-Tracker Red (Invitrogen) in pre-warmed HBSS at 37°C for 30 min, washed with HBSS, and fixed with 4% paraformaldehyde in PBS for 10 min at room temperature. Nuclei were counterstained with DAPI (1:1,000 in PBS) for 10 min. images were obtained using appropriate filter sets on a Nikon A1R+ confocal microscope (Nikon Instruments) at the Biochemistry Optical Core (BOC) Facility at UW–Madison.

### Intracellular ATP measurement

Intracellular ATP levels were measured by CellTiter-Glo^®^ Luminescent Cell Viability Assay (Promega) according to the manufacturer’s instructions. In a 96-well plate, 100 µL of CellTiter-Glo reagent was added to 100 µL of culture medium in each well. Plates were shaken for 2 min to induce cell lysis and incubated at room temperature for 10 min to stabilize the luminescent signal. Luminescence was measured using a Varioskan™ LUX Multimode Microplate Reader (Thermo Scientific)

### Mitochondrial superoxide measurement

Cells were treated with 500 nM MitoSOX™ Red Mitochondrial Superoxide Indicator (Invitrogen) prepared in HBSS with Ca^2+^ and Mg^2+^. Cells were incubated for 30 min at 37°C and 5% CO_2_. Nuclei were counterstained with Hoechst 33342 (1:2,000 in HBSS). Cells were washed three times with HBSS. Live images were taken immediately after staining. Images were obtained using appropriate filter sets on a Nikon A1R+ confocal microscope (Nikon Instruments) at the Biochemistry Optical Core (BOC) Facility at UW–Madison. Fluorescence intensity was measured using ImageJ. Where indicated, cells were treated with rotenone (250 nM) for the final 19 h of drug treatment to induce mitochondrial ROS production.

### Mitochondrial respiration analysis

Mitochondrial respiration was assessed using the Seahorse XFe24 Extracellular Flux Analyzer (Agilent) in the Rhoades lab at UW-Madison. HT29 cells (10,000 cells/well) were seeded in Seahorse XF24 cell culture microplates. Prior to the assay, the sensor cartridge was hydrated overnight in XF calibrant at 37°C in a non-CO_2_ incubator according to the manufacturer’s instructions. On the day of assay, assay medium was prepared using Seahorse XF DMEM supplemented with 100 µM pyruvate, 200 µM glutamine, and 25 mM glucose (pH 7.4 adjusted with NaOH), and warmed to 37°C before use. Cells were washed twice with 200 µL of warmed assay medium and incubated in 500 µL of assay medium for 45–60 min at 37°C in a non-CO_2_ incubator prior to measurement. The following compounds were sequentially injected: oligomycin (1.5 µM), FCCP (4 µM), and rotenone (2 µM) plus antimycin A (4 µM). Oxygen consumption rate (OCR) was measured under basal conditions and after each injection. Data were analyzed using Wave software (Agilent). Respiration values were normalized to mitochondrial content as assessed by MitoTracker staining.

### qRT-PCR

Total RNA was isolated from HT29 cells using TRIzol reagent (Invitrogen) according to the manufacturer’s instructions. cDNA was synthesized from 2 µg of total RNA using the Maxima First Strand cDNA Synthesis Kit with DNase treatment (Thermo Scientific). Quantitative PCR was performed using LightCycler 480 SYBR Green I Master Mix (Roche) on a QuantStudio™ 5 instrument (Thermo Fischer Scientific). Primers were used at a final concentration of 200 nM and the primer sequences are listed in Table S2. The thermal cycling conditions were as follows: initial denaturation at 95°C for 5 min, followed by 40 cycles of 95°C for 10 s, 60°C for 20 s, and 72°C for 15 s. A melt curve analysis was performed with 95°C for 5 s, 65°C for 1 min, and a continuous temperature ramp from 65°C to 97°C (0.075°C/s) to verify amplification specificity. Relative gene expression was calculated using the 2^-ΔΔCt^ method and normalized to *GAPDH*.

### Mitochondrial network analysis (MiNA)

After MitoTracker staining and fixation, HT29 cells were imaged using a confocal microscope with a 100x objective under Nyquist sampling conditions. Z-stack images were acquired at 0.2 µm intervals (three optical sections per cell), and maximum intensity projections were generated for analysis. Images were preprocessed and analyzed using MiNA macro, modified from Valente et al.^124^ We modified the original MiNA macro to use the Weka Trainable Segmentation^125^ Fiji plugin for improved accuracy in the segmentation of mitochondria. Mitochondrial statistics such as branching number and network size were then normalized to cytoplasm area to account for variation in cell size. Morphological parameters including mitochondrial footprint, number of individual mitochondria, number of networks, mean branch length, and mean number of branches per network were quantified.

### Cyto-ID autophagy detection and MitoTracker co-staining

Autophagy flux was assessed using the Cyto-ID^®^ Autophagy Detection Kit (Enzo Life Sciences). HT29 cells were seeded in 96-well plates and treated with chloroquine (30 µM) for 24 h prior to Cyto-ID staining to inhibit lysosomal degradation. On the day of staining, the Microplate Dual Detection Reagent was prepared by mixing 10 µL of Cyto-ID Green Detection Reagent and 10 µL of Hoechst 33342 in 10 mL of 1x Assay Buffer supplemented with 5% FBS (GenClone). For mitochondrial co-staining, 100 nM MitoTracker™ Red CMXRos (Invitrogen) was added to the staining solution. Cells were incubated with 100 µL of staining solution per well at 37°C for 30 min, washed once with 1x Assay Buffer, and imaged live using appropriate filter sets on a Nikon A1R+ confocal microscope (Nikon Instruments) at the Biochemistry Optical Core (BOC) Facility at UW–Madison.

Colocalization analysis between Cyto-ID and MitoTracker signals was performed in ImageJ using a custom macro (see the Data and code availability section) that quantified individual and overlapping fluorescence areas. Cyto-ID-mitochondria colocalization was quantified as the overlapping area normalized to the total MitoTracker area.

### *In silico* molecular docking

Molecular docking simulations were performed using DiffDock-L on the Neurosnap web platform. The crystal structure of human Folate Receptor Alpha in complex with folic acid (PDB ID: 4LRH) was obtained from the Protein Data Bank and used as the receptor structure. Ligand structures in SMILES format were submitted to DiffDock-L using default parameters. Structural visualization and figure rendering were performed using PyMOL v3.1.6.1.

### Folate receptor binding assay

Folate receptor binding assays were performed using membrane fractions isolated from HT29 cells and folic acid-d2 as a tracer ligand. The protocol was developed based on Kornilova et al.^126^ For the membrane preparation, HT29 cells were grown to confluence in three T175 flasks. Cells were placed on ice for 3 min and washed twice with ice-cold EBSS. Surface-bound folates were removed by washing cells for 60 sec on ice with acidified EBSS (pH 3.5, adjusted with acetic acid), followed by two washes with ice-cold EBSS (pH 7.4). Cells were scraped into ice-cold EBSS and centrifuged at 1,000 x g for 10 min at 4°C. Cell pellets were resuspended in 10 volumes of 10 mM Tris-HCl (pH 8.0) containing complete protease inhibitor cocktail (Roche) and homogenized for 1 min using a Bead Ruptor Elite (Omni International). The homogenate was centrifuged at 3,000 x g for 15 min at 4°C. Supernatants were collected and ultracentrifuged at 40,000 x g for 1 h at 4°C. Pellets were resuspended in suspension buffer (50 mM Tris-HCl pH 7.4, 150 mM NaCl, 25 mM n-octyl-β-D-glucopyranoside, 5 mM EDTA, 0.02% sodium azide, and protease inhibitor in sterile water), vortexed, and ultracentrifuged again at 40,000 x g for 1 h at 4°C. Membrane pellets were resuspended, protein concentration was determined and stored at −80°C for binding assay.

Binding reactions were performed in 24-well plates. Membranes (12 µg protein per well) were prepared in 350 µL of assay buffer (50 mM HEPES pH 7.2, 150 mM NaCl, 25 mM n-octyl-β-D-glucopyranoside, 5 mM EDTA, 0.02% sodium azide in sterile water) and mixed with 100 µL of 5x reaction buffer (50 mM Na2PO4, 9 mM KH2PO4, 2.5 M NaCl, 13.5 mM KCl, and 125 mM n-octyl-β-D-glucopyranoside in sterile water). After incubation for 15 min at room temperature with gentle agitation, 25 µL of competitors, folic acid-d2, were added to achieve the desired final concentrations. Samples were incubated for 2 h at room temperature with agitation. Samples were purified by solid-phase extraction (SPE) (see the ‘Metabolite extraction and LC-MS analysis’ for details) dried in a SpeedVac (Thermo Scientific), reconstituted in 40 µL of folate extraction buffer (FEB; 20 mM ascorbic acid, 20 mM ammonium acetate, 20 mM β-mercaptoethanol (BME), pH 8.1 adjusted with NaOH), and analyzed by LC-MS.

### STAT3 reporter assay

HT29 cells were seeded in 48-well plates and transfected at 70–80% confluence with the STAT3 Reporter Kit plasmids (BPS Bioscience, #79730) using Lipofectamine 3000 (Invitrogen, #L3000150) according to the manufacturer’s instructions. A total of 200 ng of STAT3 reporter plasmid was transfected per well. 24 h post-transfection, cells were treated with TNFα (10 ng/mL) for 24 h, followed by co-treatment with DAMPA or MTX in the continued presence of TNFα for an additional 24 h. Luciferase activity was measured using the Dual-Luciferase® Reporter Assay System (Promega, #E1910) according to the manufacturer’s instructions. Luminescence was measured using a Varioskan™ LUX Multimode Microplate Reader (Thermo Scientific). Firefly luciferase activity was normalized to Renilla luciferase activity.

### H&E staining

Formalin-fixed paraffin-embedded colon tissues were sectioned at 2-5 µm thickness and dried at 65°C for 25 min prior to staining. Sections were processed using the Leica ST5010 Autostainer XL (Leica Biosystems) according to the facility protocol. Sections were sequentially treated with xylene, graded ethanol, Hematoxylin MX 560, Define MX, Blue Buffer 8, and Eosin 515 Trichrome, followed by dehydration through ethanol and clearing in xylene. Images were acquired under Zeiss PrimoVert light microscope (Zeiss) by an independent investigator, coded prior to analysis, and analyzed in a blinded manner.

### Histological analysis

Four to six well-oriented crypts per mouse were analyzed. All analyses were performed in a blinded manner. Crypt length was measured using QuPath software v0.6.0 as the distance from the crypt base to the luminal surface. Goblet cells were counted per crypt based on characteristic morphology in H&E-stained sections. Neutrophil infiltrations were quantified using ImageJ, normalized to crypt area, and analyzed from the same crypt regions. Thresholds were adjusted based on visual inspection to identify positively stained cells. Clustered cells were separated using the watershed function. Neutrophils were quantified using the “Analyze Particles” function with a size threshold of 200–25,000 pixels and circularity range of 0.40–1.00.

### Serum cytokine measurement

LEGENDplex™ Mouse Anti-Virus Response Panel (13-plex) with V-Bottom Plate (BioLegend, #740622) was used to measure serum cytokine levels. For the standards, 25 µL of Matrix A was added into an assay plate, followed by adding 25 µL of serially diluted standard solutions. For serum samples, 25 µL of assay buffer was added into the assay plate, followed by adding 25 µL of 4-8x diluted serum samples. 25 µL of mixed beads were added into each well and the plate was shaken at 3,000 x g at room temperature for 2 h. After aspirating and washing the plate, 25 µL of detection antibodies were added into the plate followed by shaking at 3,000 x g at room temperature for 1 h. Then, 25 µL of SA-PE was added to each well, followed by shaking at 3,000 x g for 30 min at room temperature. After aspirating and washing the plate, the beads in the plate were resuspended in 150 µL of 1X wash buffer by pipetting and the plate was read on Attune NxT Flow Cytometer at the UW-Madison Carbone Cancer Center (UWCCC) Flow Cytometry Laboratory. Data were analyzed using the LEGENDplex™ Data Analysis Software (BioLegend). Bead populations were distinguished by FSC/SSC and internal dye intensity measured in the APC (670 nm) channel, and reporter signals were quantified in the PE (574 nm) channel.

### Immunocytochemistry

Cells grown on 96-well glass-bottom plates (#1.5 thickness, Cellvis, #NC0536760) were fixed in 4% formaldehyde and washed three times with PBS (5 min per wash). Cells were permeabilized and blocked in PBS containing 4% donkey serum (GeneTex, #GTX27475) and 0.3% Triton X-100 for 1 h at room temperature with gentle agitation. Cells were incubated with primary antibodies diluted in blocking solution overnight at 4°C with gentle agitation. The following day, cells were washed three times in PBS (5 min per wash) at room temperature and incubated with fluorophore-conjugated secondary antibodies diluted in blocking solution for 1 h at room temperature, protected from light. After washing with PBS, nuclei were stained with DAPI (1:3,000 in PBS) for 5 min. Cells were washed again with PBS before imaging. Images were acquired using appropriate filter sets on a Nikon A1R+ confocal microscope (Nikon Instruments) at the Biochemistry Optical Core (BOC) Facility at UW–Madison. Primary antibodies used for immunocytochemistry fluorescence staining are: PINK1 (1:100; Invitrogen), phospho-STAT3 (Tyr705) (pY-STAT3) (1:500, GeneTex), and phospho-STAT3 (Ser727) (pS-STAT3) (1:200, Cell Signaling). Goat anti-rabbit IgG (H+L) Cross-Adsorbed Alexa Fluor™ 488 (1:500, Invitrogen) was used as a secondary antibody.

### Fecal LCN-2 ELISA

Fecal lipocalin-2 (LCN-2) levels were quantified using a Mouse Lipocalin-2/NGAL DuoSet ELISA kit (R&D Systems, #DY1857) according to the manufacturer’s instructions. Briefly, high-binding 96-well plates (Corning, #3590) were coated overnight at room temperature with capture antibody diluted in PBS. Plates were washed with PBS containing 0.05% Tween-20 and blocked with 1% BSA in PBS for 1 h at room temperature. Fecal samples (∼20 mg) were homogenized in 500 μL PBS containing 0.1% Tween-20, centrifuged (18,200 x g, 10 min, 4°C), and the supernatant was collected for analysis. Supernatants were diluted 25x in reagent diluent. Samples and standards were added to the plate and incubated for 2 h at room temperature. After washing, detection antibody and streptavidin-HRP were sequentially applied with intermediate washing steps. Signal was developed using TMB substrate (R&D Systems, #DY999B) and stopped with stop solution (R&D Systems, #DY994). Absorbance was measured at 450 nm with wavelength correction at 540 nm. Concentrations were calculated from the standard curve.

### Graphics

The graphical abstract and schematic illustrations were created using BioRender. Chemical structures were generated using ChemDraw. Molecular docking structures and structural visualizations were rendered using PyMOL.

### Statistical analysis

Data are presented as mean ± SEM for bar graphs and scatter dot plots, unless otherwise specified. Violin plots depict data distribution, with the central line indicating the median and dotted lines representing the interquartile range (25th–75th percentiles). Each data point represents an individual biological replicate, unless otherwise specified, as defined in the figure legends. For comparisons between two groups, an unpaired two-tailed Welch’s t test was used. For non-normally distributed data, the Mann–Whitney test was applied. For comparisons among three or more groups, one-way ANOVA followed by appropriate post hoc multiple comparison tests was used. For experiments involving two independent variables, two-way ANOVA was performed. When normality assumptions were not met, the Kruskal–Wallis test was used. For the proportion of mitochondrial morphologies, the chi-square test was performed. A *p* value < 0.05 was considered statistically significant. Exact *p* values are reported in the figures.

