## Supplementary information for "Microbial metabolism of methotrexate produces a STAT3 signaling molecule that alleviates gut inflammation"

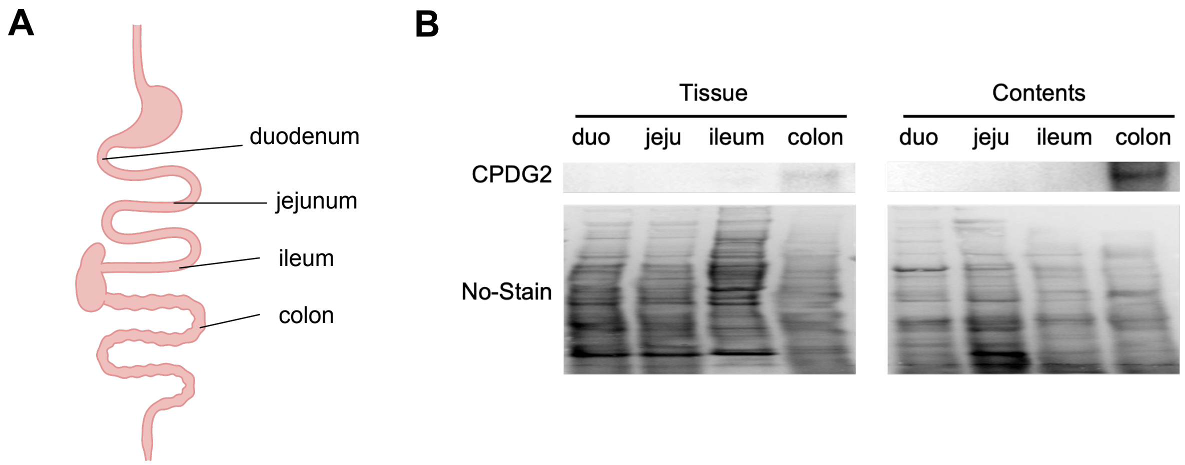


**Figure S1. CPDG2 is enriched in the distal gastrointestinal tract. Related to Figure 1.**

(A) Schematic of mouse gastrointestinal tract.

(B) Immunoblot analysis of CPDG2 in the intestinal tissues and contents from four distinct gastrointestinal segments of a healthy C57BL/6J mouse (n = 1). No-Stain was used for total protein labeling.

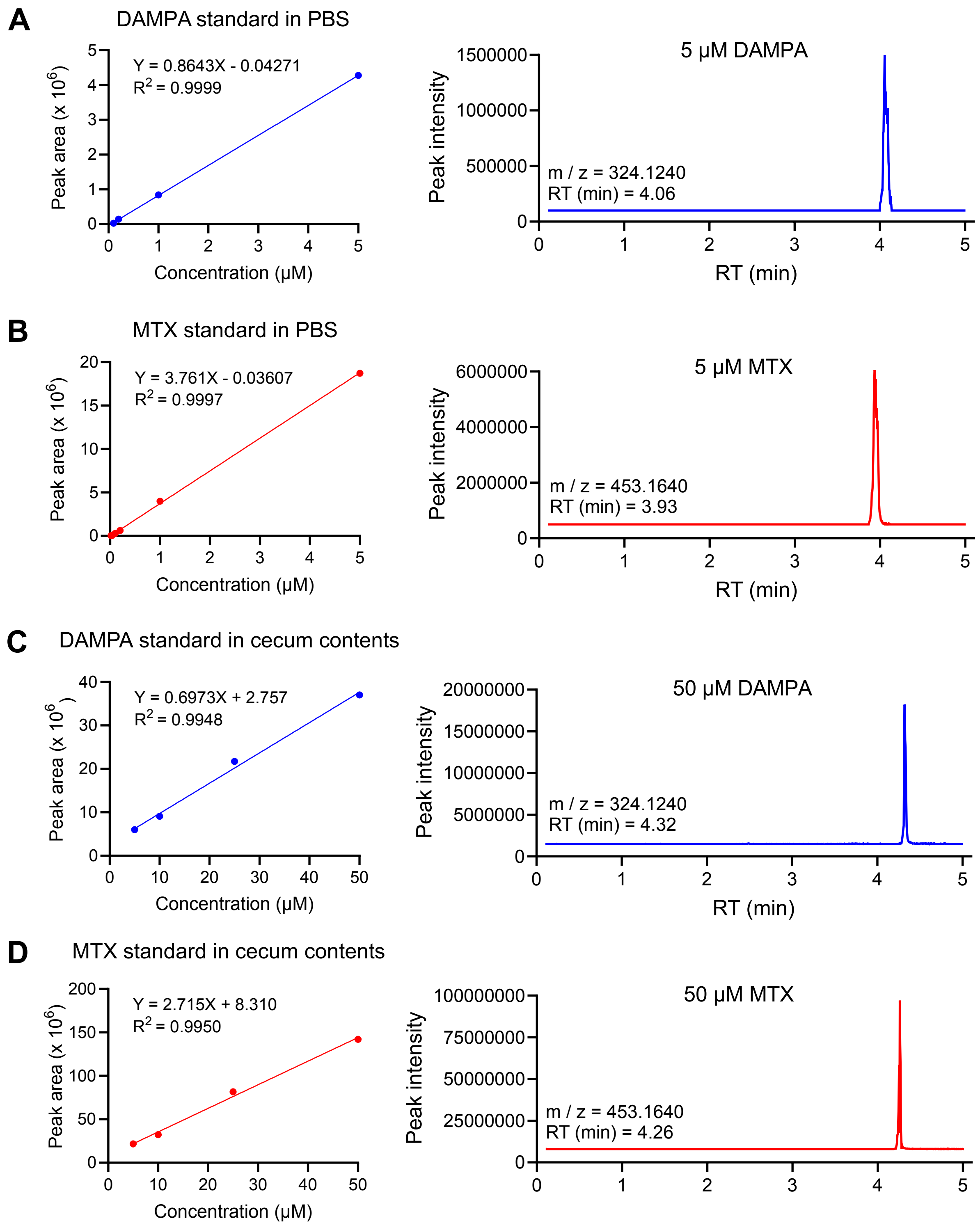


**Figure S2. Standard curves for DAMPA and MTX measured by LC-MS. Related to Figure 1.**

(A) DAMPA standard curve in phosphate-buffered saline (PBS) and representative extracted ion chromatograms at 5 μM.

(B) MTX standard curve in PBS and representative extracted ion chromatograms at 5 μM.

(C) DAMPA standard curve in mouse cecum contents and representative extracted ion chromatograms at 50 μM.

(D) MTX standard curve in mouse cecum contents and representative extracted ion chromatograms at 50 μM.

Standard curves were generated by linear regression of peak area.

m / z; mass-to-charge ratio, RT; retention time.

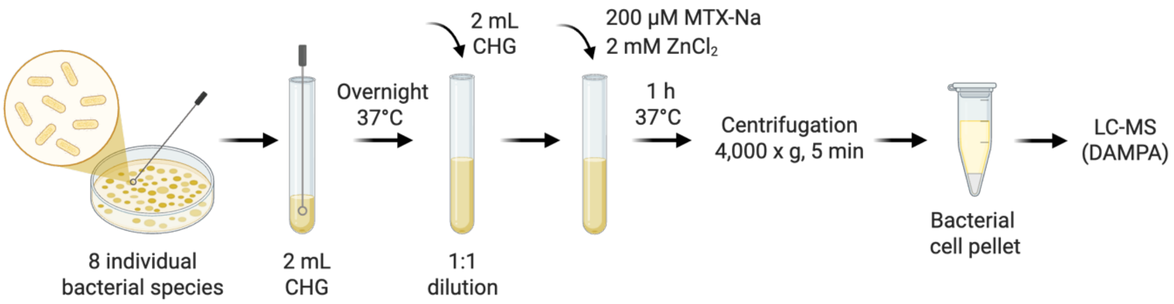


**Figure S3. Experimental scheme for *in vitro* bacterial cultures. Related to Figure 2.**

Schematic of *in vitro* anaerobic culture of individual bacterial species in Cullen–Haiser Gut (CHG) medium supplemented with MTX-Na (200 µM) and $ZnCl_2$ (2 mM) for 1 h.

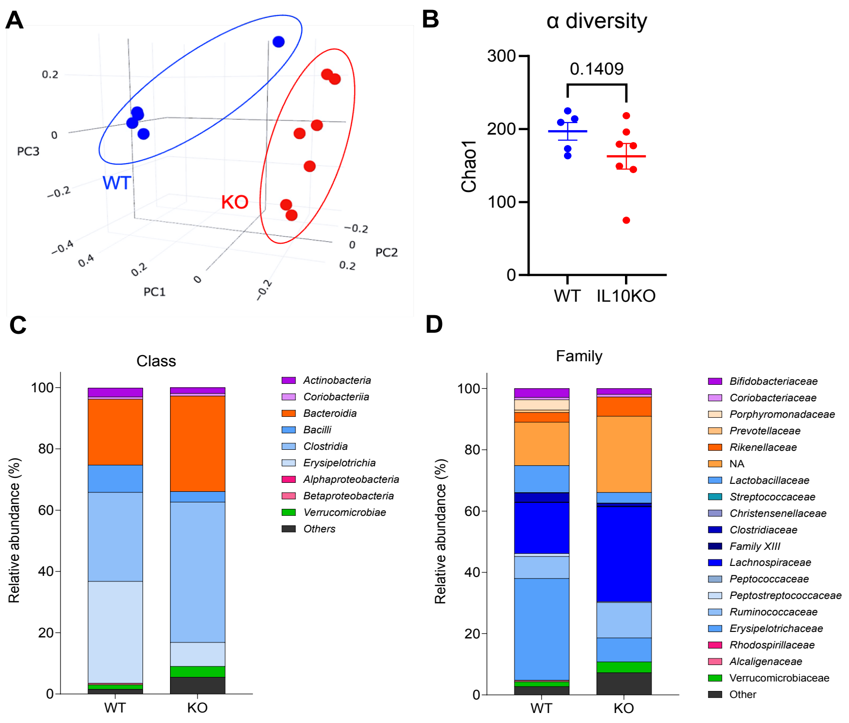


**Figure S4. Distinct gut microbial composition in WT and IL10KO mice. Related to Figure 2.**

(A) Principal coordinate analysis (PCoA) plot of fecal microbiota from WT and IL10KO mice shown in three dimensions (WT, n = 5; IL10KO, n = 7).

(B) Chao1 alpha diversity index of fecal microbiota from WT and IL10KO mice (WT, n = 5; IL10KO, n = 7, two-tailed Welch's t test).

(C and D) Relative abundance of fecal microbiota of WT and IL10KO mice at the class (C) and family (D) levels. The bar blots show the average taxonomic composition (WT, n = 5; IL10KO, n = 7).

All scatter dot plots are represented as mean ± SEM, with each data point representing an individual biological replicate. Exact *p* values are indicated in each graph.

**Table S1. Abundance of indicated phyla in PRISM and HMP2 cohorts.**

| Phylum | Healthy Relative Abundance (Control) | Direction in IBD | Estimated % Shift (IBD vs. Control) |
|---|---|---|---|
| **Firmicutes** | **55% – 75%** | **Decrease** | **-25% to -50%** |
| **Bacteroidetes** | 15% – 30% | Variable | -10% to +10% |
| **Proteobacteria** | 1% – 5% | **Increase** | **+200% to +500%** |
| **Actinobacteria** | 1% – 5% | Variable | +/-20% |
| **Verrucomicrobia** | 0.5% – 3% | **Decrease** | **-50% to -80%** |

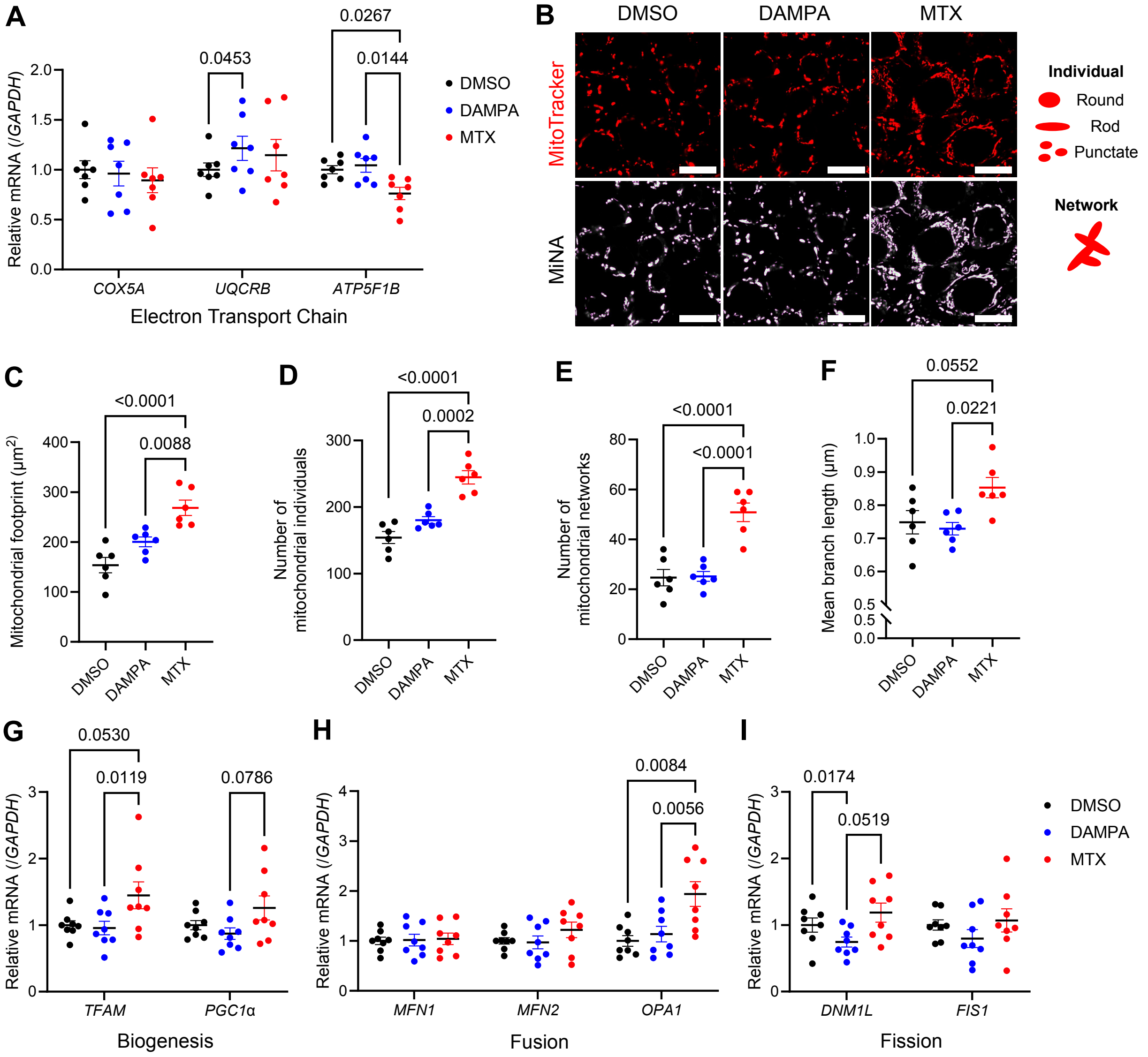


**Figure S5. MTX and DAMPA differentially regulate mitochondrial function and dynamics. Related to Figure 3.**

(A) Relative mRNA expression of genes associated with electron transport chain (Cytochrome C Oxidase Subunit 5A (*COX5A*), Ubiquinol-Cytochrome C Oxidoreductase Binding Protein (*UQCRB*), and ATP Synthase F1 Subunit Beta (*ATP5F1B*)). (n = 7 per group, two-way ANOVA). (B-F) Mitochondrial Network Analysis (MiNA) performed using ImageJ (n = 6 per group, one-way ANOVA). (B) Representative images of MitoTracker staining and corresponding MiNA outputs showing mitochondrial footprint (outlined in purple) and mitochondrial network skeletons (shown in green). Scale bar, 15 μm. Schematics illustrate the definitions of mitochondrial individuals and networks used in MiNA analysis. (C) Mitochondrial footprint ($\mu m^2$). (D) The number of mitochondrial individuals per image. (E) The number of mitochondrial networks per image. (F) Mean branch length (μm).

(G-I) Relative mRNA expression of genes associated with mitochondrial biogenesis and network dynamics, including (G) biogenesis (Mitochondrial transcription factor A (*TFAM*), and Peroxisome proliferator-activated receptor gamma coactivator 1-alpha (*PGC1α*), (H) fusion (Mitofusin-1 *(MFN1*), Mitofusin-2 (*MFN2*), and Optic Atrophy 1 (*OPA1*)), and (I) fission (Dynamin-1-like (*DNM1L*), and Mitochondria Fission 1 Protein (*FIS1*)). Expression levels were normalized to Glyceraldehyde 3-phosphate dehydrogenase (*GAPDH*) (n = 8 per group, two-way ANOVA).

All scatter dot plots are represented as mean ± SEM, with each data point representing an individual biological replicate. Exact *p* values are indicated in each graph.

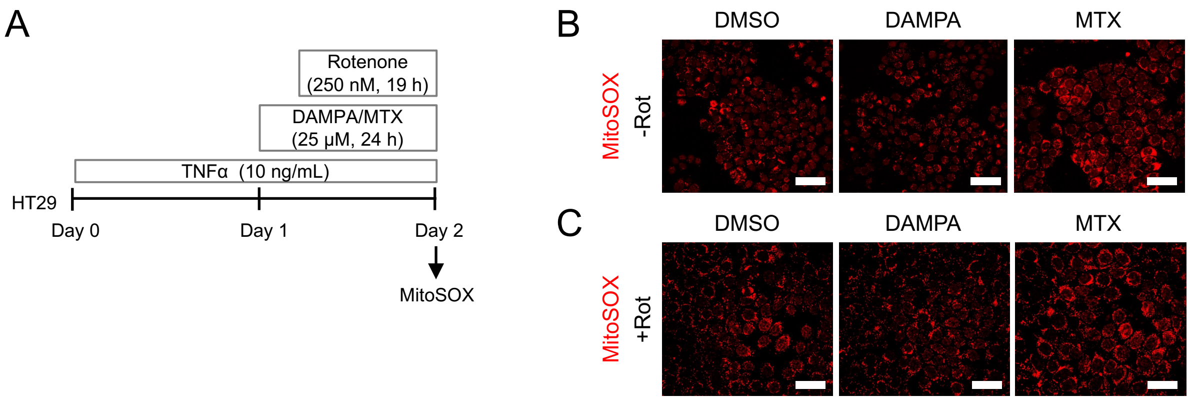


**Figure S6. MTX enhances mitochondrial superoxide levels under inflammatory conditions. Related to Figure 3.**

(A) Schematic of the experimental workflow for mitochondrial superoxide measurement using MitoSOX in HT29 cells. Cells were treated with MTX (25 µM) or DAMPA (25 µM) for 24 h under inflammatory conditions induced by TNFα (10 ng/mL), with or without rotenone (250 nM) during the final 19 h of treatment.

(B and C) Representative images of MitoSOX staining in HT29 cells treated with MTX or DAMPA, without (B) or with (C) rotenone. Scale bar, 40 µm.

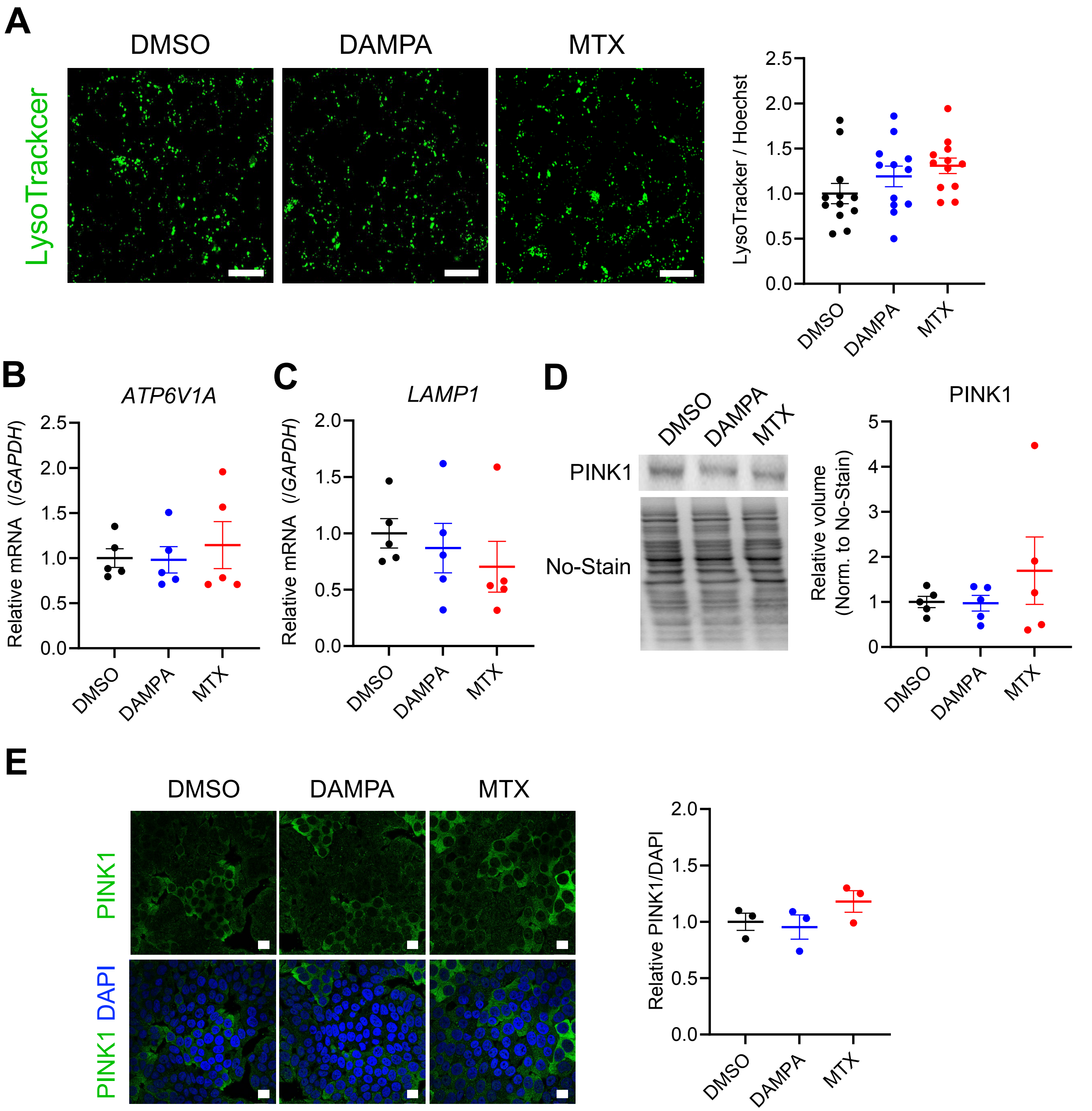


**Figure S7. DAMPA-induced autophagy occurs independently of Parkin-dependent mitophagy. Related to Figure 4.**

(A) Lysosome staining using LysoTracker in HT29 cells treated with DAMPA or MTX under TNFα-induced inflammatory conditions. Scale bar, 20 μm. LysoTracker fluorescence intensity was quantified and normalized to Hoechst 33342 (n = 12 per group).

(B and C) Relative mRNA expression of lysosomal markers, including (B) the lysosomal proton pump subunit (ATPase H+ Transporting V1 Subunit A (*ATP6V1A*)) and (C) the lysosomal membrane protein (Lysosomal Associated Membrane Protein 1 (*LAMP1*)). *GAPDH* was used for normalization (n = 5 per group).

(D and E) Abundance of the PTEN Induced Kinase 1 (PINK1) assessed by (D) western blot (n = 5) and (E) immunocytochemistry-immunofluorescence staining (n = 3 per group). Scale bar, 20 μm.

All scatter dot plots are represented as mean ± SEM, with each data point representing an individual biological replicate.

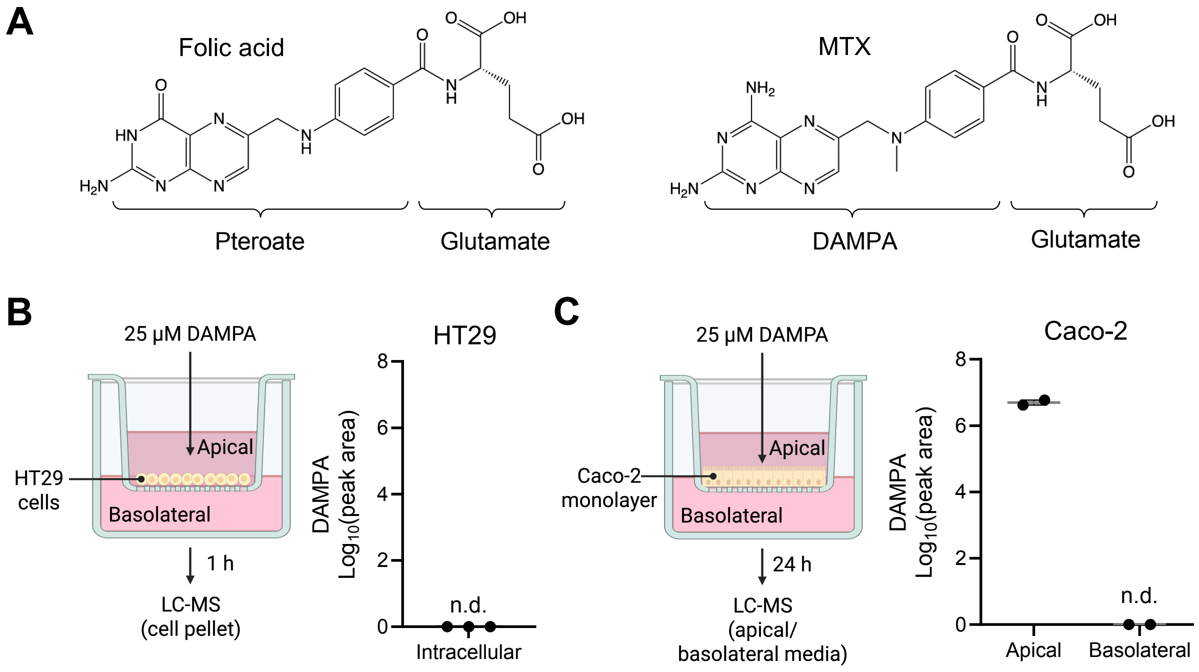


**Figure S8. DAMPA does not enter intestinal epithelial cells. Related to Figure 5.**

(A) Chemical structure of folic acid, composed of pteroate moiety and glutamate, and MTX, composed of DAMPA and glutamate.

(B) Apical DAMPA treatment in HT29 cells cultured on Transwell inserts. Cells were treated apically with DAMPA (25 µM) for 1 h, followed by DAMPA quantification in cell pellets by LC-MS (n.d. = not detected, n = 3).

(C) Apical DAMPA treatment in differentiated Caco-2 monolayers cultured on Transwell inserts. Cells were treated apically with DAMPA (25 µM) for 24 h, and apical and basolateral medium were collected for DAMPA measurement by LC-MS (n.d. = not detected, n = 2).

All scatter dot plots are represented as mean ± SEM, with each data point representing an individual biological replicate.

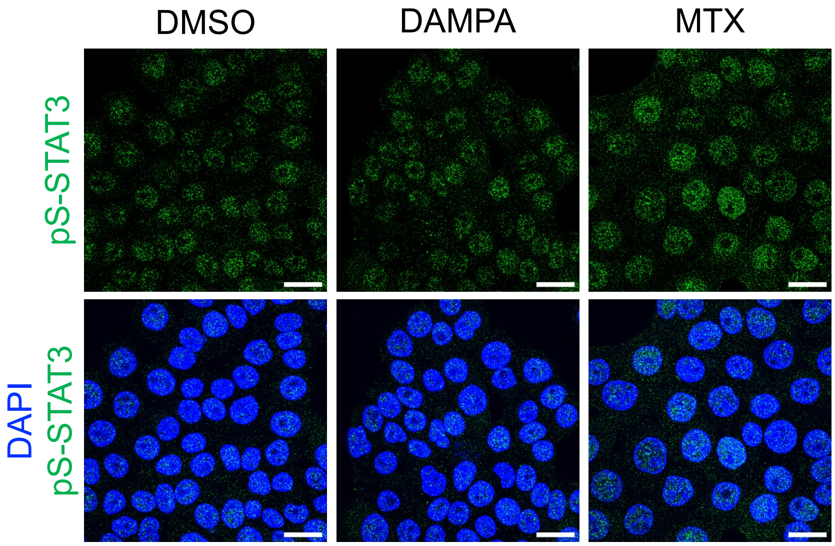


**Figure S9. Phosphorylated STAT3 at Ser727 is predominantly localized to the nucleus. Related to Figure 5.**

Immunofluorescence staining of phosphorylated STAT3 at Ser727 (pS-STAT3) in HT29 cells treated with TNFα (10 ng/mL) followed by DAMPA (25 μM) or MTX (25 μM) treatment for 24 h. Nuclei were counterstained with DAPI. Scale bar, 20 μm.

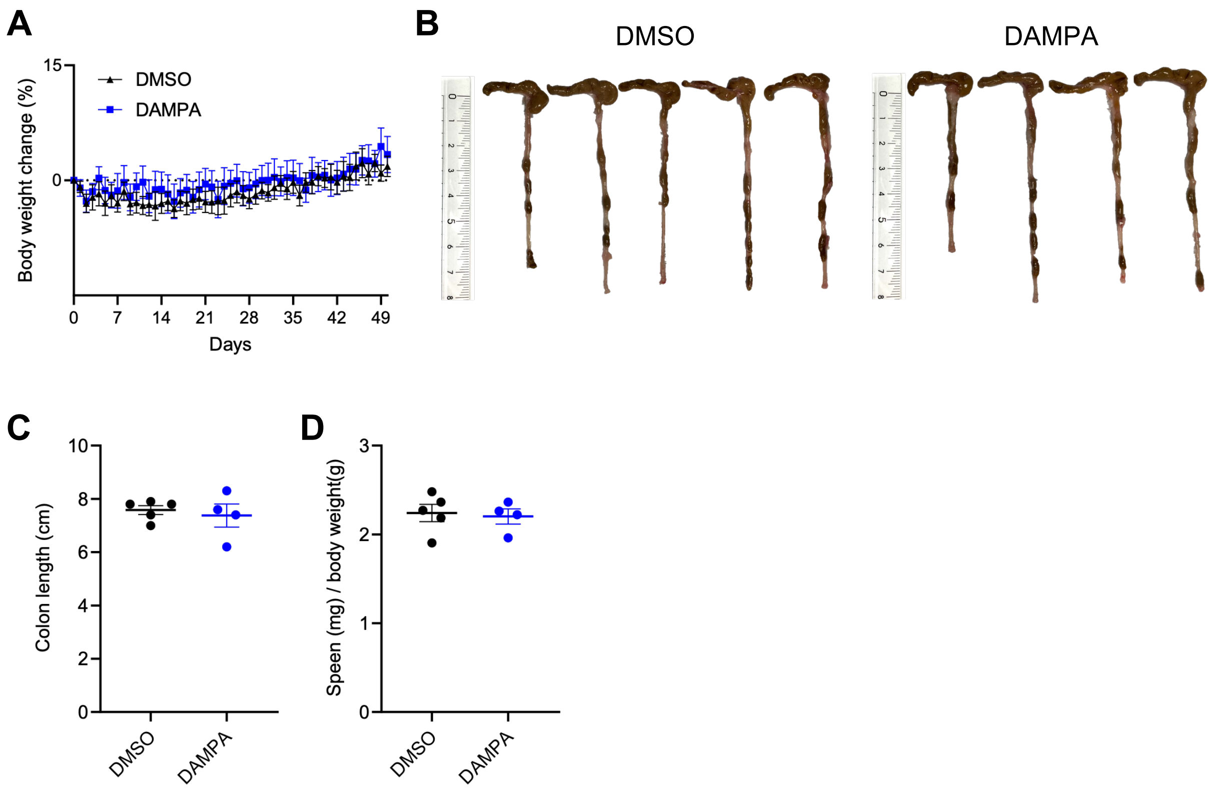


**Figure S10. DAMPA does not affect colon length or spleen weight in IL10KO mice. Related to Figure 6.**

(A) Body weight changes during 50 days of daily oral DAMPA administration. Data are shown as mean ± SEM.

(B and C) Colon images and length measurement at the time of sacrifice. Length scale in centimeters.

(D) Spleen weight normalized to body weight.

(C and D) Data are represented as mean ± SEM, with each data point representing an individual biological replicate.

**Table S2. qPCR primer sequences.**

| Gene | Forward (5' - 3') | Reverse (5' - 3') |
|---|---|---|
| COX5A | TGATATGGTTCCAGAGCCCA | CCTGGATGACATAGGGGTAGA |
| UQCRB | AGTGGCTGGATGGTATTCGA | CAGGTCCAGTGCCCTCTTAA |
| ATP5F1B | TGCATTATTGGGCCGAATCC | CAGTCAAGTCATCAGCAGGC |
| PGC1α | GAGAGTCTGAGAGGGCCAAG | CCTCAGTTCTGTCCGTGTTG |
| TFAM | AGAAGAATTGCCCAGCGTTG | CTGACTTGGAGTTAGCTGTTCT |
| MFN1 | GGATTGGCGTCCGTTACATC | TGGTCCAGCTCAGTCTTTCA |
| MFN2 | TACACTGGCTCCAACTGCAG | TTTCTTGTTCATGGCGGCAA |
| OPA1 | TGGAAGAATCGGACCCAAGA | ACTCCTCGGGATTCAAGGTT |
| DNM1L | GGAACAGCGAGATTGTGAGG | TGGCCTACTAGCTCACTCTG |
| FIS1 | CAGTTTGAGTACGCCTGGTG | TGTTCCTCCTTGCTCCCTTT |
| GAPDH | GACAGTCAGCCGCATCTTCT | GCGCCCAATACGACCAAATC |
| ATP6V1A | TGCGTGCCTTGGATGAATAC | AGTTTTGCTACCTCCAGAGTGA |
| LAMP1 | TGACCGTAACGCTCCATGAT | GCTCACGTTGTACTTGTCCA |

All primers are designed for human gene expression analysis
